# Elimination of subtelomeric repeat sequences exerts little effect on telomere essential functions in *Saccharomyces cerevisiae*

**DOI:** 10.1101/2023.08.29.555450

**Authors:** Can Hu, Xue-Ting Zhu, Ming-Hong He, Yangyang Shao, Zhongjun Qin, Zhi-Jing Wu, Jin-Qiu Zhou

## Abstract

Telomeres, which are chromosomal end structures, play a crucial role in maintaining genome stability and integrity in eukaryotes. In the baker’s yeast *Saccharomyces cerevisiae*, the X-and Y’-elements are subtelomeric repetitive sequences found in all thirty-two and seventeen telomeres, respectively. While the Y’-elements serve as a backup for telomere functions in cells lacking telomerase, the function of the X-elements remains unclear. This study utilized the *S. cerevisiae* strain SY12, which has three chromosomes and six telomeres, to investigate the role of X-elements (as well as Y’-elements) in telomere maintenance. Deletion of Y’-elements (SY12^YΔ^), X-elements (SY12^XYΔ+Y^), or both X- and Y’-elements (SY12^XYΔ^) did not impact the length of the terminal TG_1-3_ tracks or telomere silencing. However, inactivation of telomerase in SY12^YΔ^, SY12^XYΔ+Y^, and SY12^XYΔ^ cells resulted in cellular senescence and the generation of survivors. These survivors either maintained their telomeres through homologous recombination-dependent TG_1-3_ track elongation or underwent microhomology-mediated intra-chromosomal end-to-end joining. Our findings indicate the non-essential role of subtelomeric X-and Y’-elements in telomere regulation in both telomerase-proficient and telomerase-null cells and suggest that these elements may represent remnants of *S. cerevisiae* genome evolution. Furthermore, strains with fewer or no subtelomeric elements exhibit more concise telomere structures and offer potential models for future studies in telomere biology.

## INTRODUCTION

Telomeres, specialized nucleoprotein structures located at the end of linear chromosomes in eukaryotic cells, are crucial for maintaining genomic stability and protecting chromosomal ends from being perceived as DNA breaks (Wellinger and Zakian, 2012). In the budding yeast *Saccharomyces cerevisiae*, telomeric DNA consists of approximately ∼300±75 base pairs of C_1-3_A/TG_1-3_ repeats with a 3’ G-rich single-stranded overhang (Wellinger and Zakian, 2012). Adjacent to the telomeric TG_1-3_ repeats, there are subtelomeric repeat elements known as X- and Y’-elements, which vary between telomeres, as well as strains (Chan and Tye, 1983a, b; Louis, 1995). The Y’-elements, immediately internal to the telomeric repeats, are present as a tandem array of zero to four copies, they fall into two major size classes, 6.7 kb Y’-long (Y’-L) and 5.2 kb Y’-short (Y’-S) (Chan and Tye, 1983a, b). Y’-elements are highly conserved with only ∼2% divergence between strains (Louis and Haber, 1992). One entire Y’-element contains two large open reading frames (ORFs), an ARS consensus sequence (ACS) and a STAR element (Subtelomeric anti-silencing regions) consisting of binding sites for Tbf1 and Reb1 (Chan and Tye, 1983a, b; Fourel et al., 1999; Louis and Haber, 1992). The X-element, a much more heterogeneous sequence abutting Y’-elements or telomeric repeats, contains the 473 bp “core X” sequence and the subtelomeric repeats (STRs) A, B, C and D (Louis and Haber, 1991; Louis et al., 1994). The STRs are found in some chromosome ends, while the “core X” sequence is shared by all chromosomes. Recent long-read sequencing shows that subtelomeric regions display high evolutionary plasticity and are rich in a various of structure variants such as reciprocal translocations, transpositions, novel insertions, deletions and duplications (O’Donnell et al., 2023).

Telomeric DNA elongation primarily relies on telomerase, an enzyme comprising a reverse transcriptase, an RNA component, and accessory factors (Palm and de Lange, 2008; Wellinger and Zakian, 2012). In *S. cerevisiae*, the telomerase holoenzyme consists of the reverse transcriptase Est2, the RNA template TLC1, and accessory factors Est1, Est3, Pop1/Pop6/Pop7 proteins (Lemieux et al., 2016; Lendvay et al., 1996; Lundblad and Szostak, 1989; Singer and Gottschling, 1994). In the absence of telomerase, homologous recombination can take place to replicate telomeres, resulting in telomerase-deficient “survivors” (Lundblad and Blackburn, 1993; Teng and Zakian, 1999). These survivors are broadly categorized into Type I and Type II based on distinct telomere structures (Lundblad and Blackburn, 1993; Teng and Zakian, 1999). Type I survivors possess tandem amplified Y’-elements (both Y’-L and Y’-S) and very short TG_1-3_ tracts, indicating that Y’-elements serve as substrates for homologous recombination. Type II survivors display long heterogeneous TG_1-3_ tracts. On solid medium, approximately 90% of the survivors are Type I, while 10% are Type II (Teng et al., 2000). However, in liquid culture, Type II survivors grow faster and eventually dominate the population (Teng and Zakian, 1999). The proteins required for Type I and Type II survivor formation appear to be different. Type I survivors depend on Rad51, Rad54, Rad55, Rad57, and Pif1 (Chen et al., 2001; Hu et al., 2013; Le et al., 1999). while the formation of Type II survivors requires the Mre11/Rad50/Xrs2 (MRX) complex, KEOPS complex, Rad59, Sgs1, and Rad6, most of which are critical for DNA resection (Chen et al., 2001; He et al., 2019; Hu et al., 2013; Johnson et al., 2001; Le et al., 1999; Louis, 2001; Nicolette et al., 2010; Teng et al., 2000; Wellinger and Zakian, 2012; Wu et al., 2017). Although Type I and Type II pathway are working independently, Kockler et al. found that the proteins involved in each pathway can work together via two sequential steps and contribute to a unified ALT (Alternative lengthening of telomeres) process (Kockler et al., 2021).

The amplification of Y’-elements represents a significant feature of telomere recombination in telomerase-null Type I survivors (Lundblad and Blackburn, 1993; Teng and Zakian, 1999), and as a result, extrachromosomal Y’ circular DNAs have been observed in Type I survivors (Larrivee and Wellinger, 2006). Additionally, Y’-element acquisition has been observed in the initiation step of pre-senescence, suggesting a potential role for Y’-elements in Type II survivor formation (Churikov et al., 2014). Furthermore, Y’-elements are mobilized through a transposition-like RNA-mediated process involving Ty1 activity in telomerase-negative survivors (Maxwell et al., 2004). Y’-elements also express potential DNA helicases, Y’-Help, in telomerase-null survivors (Yamada et al., 1998). Thus, Y’-elements play a significant role as donors in homologous recombination-mediated telomere maintenance. The functions of X-elements, on the other hand, are less clear. The “core X” sequence consists of an ACS element and, in most cases, an Abf1 binding site (Louis, 1995), and acts as a protosilencer (Lebrun et al., 2001). In contrast, STRs and Y’-STAR possess anti-silencing properties that limit the spreading of heterochromatin (Fourel et al., 1999). Interestingly, a previous study demonstrated that telomeres with Χ-only ends (containing only X-elements) were more efficiently elongated compared to those with X-Y’ ends (containing both X- and Y’-elements) in *tel1*Δ *rif1*Δ strains (Craven and Petes, 1999). Moreover, subtelomeric elements (including X-elements) and associated factors like Reb1 and Tbf1 antagonize telomere anchoring at the nuclear envelope (Hediger et al., 2006). However, considering that X-elements are present in all telomeres while Y’-elements are not, the specific functions of X- and Y’-elements in genome integrity after the evolution of telomerase have long been a subject of questioning (Jager and Philippsen, 1989; Zakian and Blanton, 1988).

In wild-type yeast strain BY4742, there are eight Y’-S and eleven Y’-L elements at the thirty-two telomere loci. Additionally, each telomere locus contains one X-element. The genetic deletion of all X- and Y’-elements to directly investigate the roles of X- and Y’-elements in genome integrity is a challenging and complex task. In this study, we utilized recently reported chromosome-fused budding yeast strains (Shao et al., 2018) to eliminate both X- and Y’-elements completely. This approach allows us to reinvestigate the roles of X-and Y’-elements at telomeres.

## RESULTS

### Telomere recombination in telomerase-null chromosome-fused yeast strains SY1 to SY12

The functions of Y’-elements have been previously linked to telomere recombination (Churikov et al., 2014; Larrivee and Wellinger, 2006; Lundblad and Blackburn, 1993; Teng and Zakian, 1999). To further investigate the role of Y’-elements in telomere recombination, we utilized a series of chromosome-fused budding yeast strains derived from the wild-type BY4742 strain, including SY1, SY3, SY5, SY7, SY8, SY9, SY10, SY11, SY12 and SY13 (also referred to as SYn for convenience) (Figure 1A) (Shao et al., 2018). The remaining subtelomeric elements in SY8 to SY13 strains are listed in supplementary file 2. We excluded SY14 from these experiments since the presence of circular chromosome was prominent in SY14 *tlc1*Δ cells (one fused chromosome) (Wu et al., 2020), We generated haploid SYn *tlc1*Δ *TLC1* strains by deleting the chromosomal copy of the *TLC1* gene and introducing a plasmid-borne wild-type *TLC1* gene (pRS316-*TLC1*). Clones that lost the pRS316-*TLC1* plasmid (containing the *URA3* marker) were identified upon counter-selection on 5’-fluoroorotic-acid (5’-FOA) plates and were subsequently re-streaked on YPD plates for at least 9 cycles for survivor formation (referred to as the “multiple-colony streaking assay” in Methods). The telomere patterns of the survivors were then determined through Southern blotting assay (Figure 1B-D).

The canonical telomerase-independent survivors can be broadly categorized into two types: Type I and Type II survivors, based on the restriction fragments generated after XhoI digestion (Lundblad and Blackburn, 1993; Teng and Zakian, 1999). Type I survivors exhibit tandem duplication of Y’-elements and very short TG_1-3_ tracts, while Type II survivors contain long heterogeneous TG_1-3_ sequences. Consistent with previous reports, BY4742 *tlc1*Δ cells generated both Type I (subtelomeric Y’-element recombination) and Type II (TG_1-3_ recombination) survivors (Figure 1B) (Hu et al., 2013). Intriguingly, as the number of chromosomes decreased, the frequency of Type II survivors gradually diminished, while Type I survivors became the predominant type (Figure 1B-D). Furthermore, non-canonical survivors with distinct patterns from Type I or Type II emerged in SY9 *tlc1*Δ (six chromosomes), SY10 *tlc1*Δ (five chromosomes), SY11 *tlc1*Δ (four chromosomes), SY12 *tlc1*Δ (three chromosomes) and SY13 *tlc1*Δ (two chromosomes) (Figure 1C and 1D indicated by triangles at the bottom of the panels). Notably, the Y’-telomere band of ∼1.2 kb was not detected in two clones of SY11 *tlc1*Δ cells (clones 2 and 5), the majority of clones of SY12 *tlc1*Δ cells (except for clones 9, 14, and 15), and the majority of clones of SY13 *tlc1*Δ cells (except for clones 1, 4, 8 and10) (Figure 1D). We speculate that either the Y’-elements have eroded or the chromosomal ends containing Y’-elements have fused with other ends in these non-canonical survivors. These findings suggest that the ratio of survivor types is influenced by the number of chromosomes.

**Figure 1.**
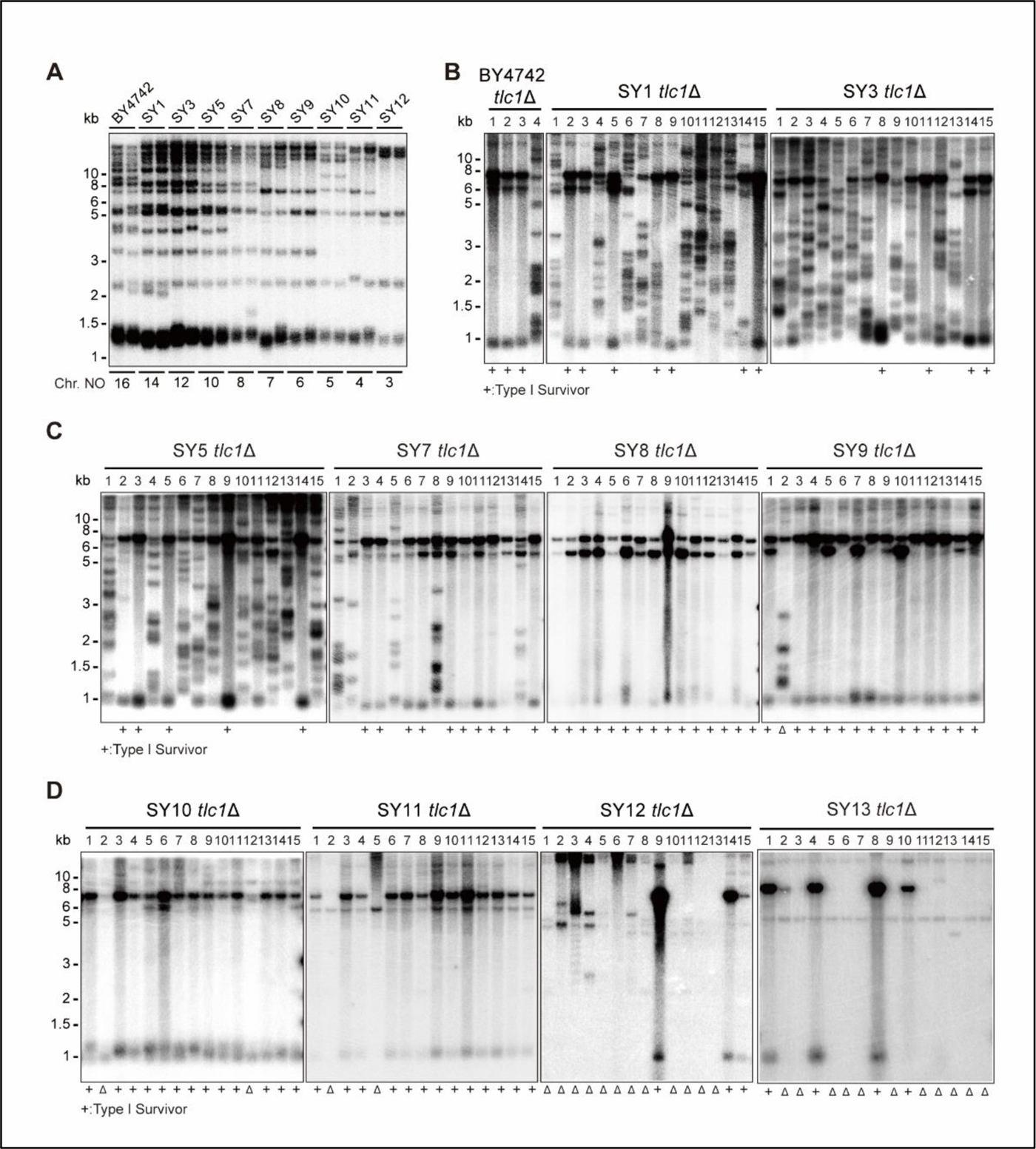
Telomere structures in SYn *tlc1*Δ survivors. Telomere Southern blotting assay was performed to examine telomere structure. The genomic DNA extracted from BY4742 (wild type) and SYn strains (labeled on top) was digested with XhoI and subjected to Southern hybridization with a TG1-3 probe. (**A**) Telomerase-proficient strains (labeled on top), whose chromosome numbers are labeled at the bottom. Two independent clones of each strain were examined. (**B-D**) SYn *tlc1*Δ survivors generated on plates. Four (BY4742 *tlc1*Δ) and fifteen (SYn *tlc1*Δ) individual survivor clones (labeled on top of each panel) of each strain were examined. “+” at the bottom indicates Type I survivors. “Δ” marks the survivors which are non-canonical Type I or Type II.

### Characterizing the survivor pattern in SY12

To determine the chromosomal end structures of the non-canonical survivors shown in Figure 1, we selected SY12 *tlc1*Δ survivors for further analysis. In the SY12 strain, there are six telomeres corresponding to the native chromosomes I-L, X-R, XIII-L, XI-R, XVI-L, and XIV-R. We employed Southern blotting after NdeI digestion to validate the telomere and subtelomere structures (Figure 2—figure supplement 1A). The results revealed that, in the SY12 strain used in our study, only the XVI-L telomere contained a single copy of the Y’-element, while all telomeres harbored X-elements (Figure 2—figure supplement 1B). For simplicity, we referred to the chromosomes containing the original I, XIII, and XVI as chromosome 1, 2, and 3, respectively (Figure 2A, left panel).

We conducted a re-examination of telomere recombination upon telomerase inactivation in SY12 cells. Deletion of *TLC1* in SY12 cells resulted in cell senescence, and different clones recovered at various time points in liquid medium (Figure 2B). Telomere Southern blotting analysis showed progressive shrinking of the telomeric XhoI fragments over time, and TG_1-3_ recombination occurred to maintain telomeres (Figure 2C). Since the liquid culture contained a mixture of different colonies, we employed a multiple-colony streaking assay and Southern blotting analysis to examine the telomere patterns of fifty independent SY12 *tlc1*Δ survivors (Figure 2D and Figure 2—figure supplement 2). Among these survivors, eight clones (labeled in red, 16% of the survivors tested) exhibited the typical Type I telomere structure characterized by Y’-element amplification (Figure 2D and 2E). This was confirmed by Southern blotting analysis using a Y’ probe (Figure 2—figure supplement 2). The emergence of Type I survivor in SY12 strain which only contain one Y’-element indicates that multiple Y’-elements in tandem are not strictly required for Type I formation. Clone 1 (labeled in orange, 2% of the survivors tested) displayed heterogeneous telomeric TG_1-3_ tracts (Figure 2D and 2E), indicating it was a Type II survivor. This was further confirmed by restoring the telomere length to the level observed in SY12 cells through the re-introduction of the *TLC1* gene into one representative clone (named SY12 *tlc1*Δ-T1) and subsequent passaging on yeast complete (YC) medium lacking uracil (Ura-) for 20 cycles (Figure 2—figure supplement 3A).

Notably, ten of the examined clones (labeled in blue, 20% of the survivors tested) displayed no telomere signals associated with canonical Type I or Type II survivors (Figure 2D and 2E). Their hybridization patterns were strikingly similar to those of SY14 *tlc1*Δ survivors (Wu et al., 2020), which survived through intra-chromosomal circularization. To investigate whether the three chromosomes in these SY12 *tlc1*Δ survivors had undergone intra-chromosomal fusions, we selected a clone, namely SY12 *tlc1*Δ-C1 and performed PCR-mapping assay to determine the erosion points of each chromosome end, as previously described (Wu et al., 2020). A PCR product of the predicted length would be obtained only if the corresponding chromosome region was intact. The PCR-mapping assay precisely identified the borders of telomere erosion for the three chromosomes in SY12 *tlc1*Δ-C1 cells. For chromosome 1 (Figure 2A, left panel), the chromosome regions approximately 3.3 kb and 1.9 kb proximal to telomere I-L and X-R, respectively, had been lost (Figure 2—figure supplement 4 and supplementary file 3). Regarding chromosome 2, the terminal ∼3.8 kb of telomere XIII-L and ∼2.5 kb of telomere XI-R remained intact (Figure 2—figure supplement 4 and supplementary file 3). For chromosome 3, the terminal ∼0.1 kb of telomere XVI-L was intact, while the terminal ∼3.4 kb of telomere XIV-R was preserved (Figure 2—figure supplement 4 and supplementary file 3). To confirm the chromosome fusion events, we performed PCR-sequencing analysis. If a given pair of primers, oriented to different chromosome ends, produced PCR products, it indicated that the corresponding arms had fused. The results revealed that the three chromosomes in SY12 *tlc1*Δ-C1 cells had undergone intra-chromosomal fusions through microhomology-mediated end joining (MMEJ) (Wu et al., 2020), resulting in the formation of circular chromosomes (Figure 2F and supplementary file 3). Notably, the fusion junctions of the three chromosomes in SY12 *tlc1*Δ-C1 cells differed in nucleotide sequence and length (22 bp, 8 bp, and 5 bp in chromosomes 1, 2, and 3, respectively). Moreover, the sequences involved in the ends-fusion were not perfectly complementary (Figure 2F). For example, the fusion sequence of chromosome 3 was 5 bp long and contained one mismatch. To further verify the chromosome structure in the “circular survivors” SY12 *tlc1*Δ-C1 (Figures 2F), we performed the pulsed-field gel electrophoresis (PFGE) analysis. Control strains included SY12 (three linear chromosomes) and SY15 (one circular chromosome). The PFGE result confirmed that like the single circular chromosome in SY15 cells, the circular chromosome in the SY12 *tlc1*Δ-C1 survivors couldn’t enter the gel, while the linear chromosomes in SY12 were separated into distinct bands, as expected (Figure 2—figure supplement 5). Thus, the survivors shown in Figure 2D, which displayed an identical hybridization pattern to the SY12 *tlc1*Δ-C1 clone, were all likely “circular survivors”. Consistently, the telomere signals detected in the SY12 strain were still not observed in the SY12 *tlc1*Δ-C1 survivor after reintroducing a plasmid-borne wild-type *TLC1* gene (Figure 2—figure supplement 3B).

Twelve clones of SY12 *tlc1*Δ survivors (labeled in green, 24% of the survivors tested) exhibited no Y’-telomere signals compared to SY12 cells but displayed different lengths of TG_1-3_ tracts (Figure 2D and 2E). Due to their non-canonical telomere structures, characterized by the absence of both Y’-amplification and superlong TG_1-3_ sequences, we designated these SY12 *tlc1*Δ survivors (labeled in green, Figure 2D) as Type X. In Type X survivors, the DNA bands with sizes of approximately 2.3 kb, 5.1 kb, 15.3 kb, 18.5 kb, and 21.9 kb were roughly comparable to the telomeres of XI-R, X-R, I-L, XIII-L, and XIV-R in SY12 cells (indicated on the left in the panel). The newly emerged band at approximately 4.3 kb likely originated from the XVI-L telomere (indicated by the red arrow on the right in the panel) (Figure 2D), where the Y’-elements had been eroded, leaving only the TG_1-3_ tracts at the very ends (Figure 2A, right panel). It remains unclear whether Y’-element erosion is common in telomerase-null BY4742 type II survivors. However, in SY12 *tlc1*Δ cells, the remaining single copy of the Y’-element couldn’t find homology sequences to repair telomeres, whereas the multicopy X-element could easily find homology sequences to repair telomeres and form the type X survivors. To verify this notion, we reintroduced the *TLC1* gene into one representative clone (named SY12 *tlc1*Δ-X1) and examined the telomere length. As expected, the telomeres of X-R and XI-R were restored to the lengths observed in wild-type SY12 cells, and accordingly, the newly emerged 4.3 kb band was also elongated (Figure 2G). Given that the restriction fragments of telomeres I-L (15.3 kb), XIII-L (18.5 kb), and XIV-R (21.9 kb) were quite long, detecting minor changes in telomere length was challenging under the assay conditions of Southern blotting. To determine the chromosomal end structure of the Type X survivor, we randomly selected a typical Type X survivor, and performed PCR-sequencing analysis. The results revealed the intact chromosome ends for I-L, X-R, XIII-L, XI-R, and XIV-R, albeit with some mismatches compared with the *S. cerevisiae* S288C genome (http://www.yeastgenome.org/), which possibly arising from recombination events that occurred during survivor formation. Notably, the sequence of the Y’-element in XVI-L could not be detected, while the X-element remained intact (Figure 2—figure supplement 6). These data indicated that Type X survivors possess linear chromosomes with telomeres terminating in TG_1-3_ repeats, while the Y’-element has been eroded (Figure 2A, right panel). Consistently, no Y’ signals were detected in these twelve Type X survivors (labeled in green, Figure 2—figure supplement 2), suggesting that the Y’-element has not been translocated to other telomeres and is not essential for yeast cell viability.

In addition to the aforementioned Type I, Type II, circular, and Type X survivors, there were some clones (labeled in black, 38% of the survivors tested) which exhibited non-uniform telomere patterns and were not characterized (Figure 2D and Figure 2E). We speculated that combinations of diverse mechanisms were occurring within each “uncharacterized survivor”. For instance, in the case of two survivors (clones 9 and 18, Figure 2D) in which only one hybridization signal could be detected, pointing to the possibility that two chromosomes underwent intra-chromosomal fusions while one retained its ends through TG_1-3_ recombination. However, the sizes of the two telomere restriction fragments on the linear chromosome were too close to be distinguished and separated, resulting in only one hybridization signal. Alternatively, it is also plausible that three chromosomes experienced intra-chromosomal fusions, with one fusion point containing TG_1-3_ repeats. For the uncharacterized clones 4, 5, 7, 15, and 43, they exhibited significant amplification of TG_1-3_ sequences, and the telomeres of these survivors did not resolve into distinct bands (Figure 2D). We hypothesize that the observed telomere patterns in these survivors could be attributed to extensive TG_1-3_ recombination. However, we cannot exclude the possibility of coexisting diverse mechanisms within a survivor, such as telomere elongation through TG_1-3_ amplification, as well as intra- and inter-chromosomal fusions. Since we couldn’t figure out the telomere structures in these survivors, we classified them as “uncharacterized survivors”.

To further determine the genetic requirements for survivors in SY12, we constructed the SY12 *tlc1*Δ *rad52*Δ pRS316-*TLC1* strain. The plasmid-borne wild-type *TLC1* gene (pRS316-*TLC1*) was counter-selected on 5’-FOA plates. SY12 *tlc1*Δ *rad52*Δ cells were measured by the cell viability assay (see Materials and methods). The results showed double deletion of *TLC1* and *RAD52* in SY12 strain could slightly accelerate senescence, and SY12 *tlc1*Δ *rad52*Δ survivors could be generated but took much longer to recover than the SY12 *tlc1*Δ survivors (Figure 2—figure supplement 7A), suggesting that Rad52 is not strictly required for survivor generation in the SY12 strain in liquid. We also passaged SY12 *tlc1*Δ *rad52*Δ cells on solid medium until survivor emerged. Southern blotting of twenty-five clones revealed that neither Type I nor Type II survivors were found, and instead circular survivors except clone 20 were obtained (labeled in blue, Figure 2—figure supplement 7B). We conclude that the formation of circular survivors in the SY12 *tlc1*Δ *rad52*Δ strain is mediated by MMEJ as observed in the SY14 *tlc1*Δ *rad52*Δ strain (Wu et al., 2020), but not *RAD52* mediate pathways. Since no Type X survivor was detected in SY12 *tlc1*Δ *rad52*Δ strain, we constructed the SY12 *tlc1*Δ *rad51*Δ pRS316-*TLC1* and SY12 *tlc1*Δ *rad50*Δ pRS316-*TLC1* strain to investigate on which pathway the formation of the Type X survivor relied. After being counter-selected on 5’-FOA plates, cells were passaged on solid medium until survivor arose. Southern blotting assay indicated the emergence of Type X survivors even in the absence of Rad51 (labeled in green, clone 2, 5, 11 and 18, Figure 2—figure supplement 8A). In contrast, no Type X survivor was detected in the SY12 *tlc1*Δ *rad50*Δ strain (Figure 2—figure supplement 8B). These data suggest that the formation of the Type X survivor depends on Rad50-mediated Type II pathway.

Taken together, our results indicate that telomerase inactivation in SY12 cells leads to cell senescence and the emergence of survivors with diverse telomere patterns, including Y’-amplification (Type I), elongated TG_1-3_ tracts (Type II), intra-chromosomal end-to-end joining (circular), Y’-loss (Type X), and uncharacterized.

**Figure 2.**
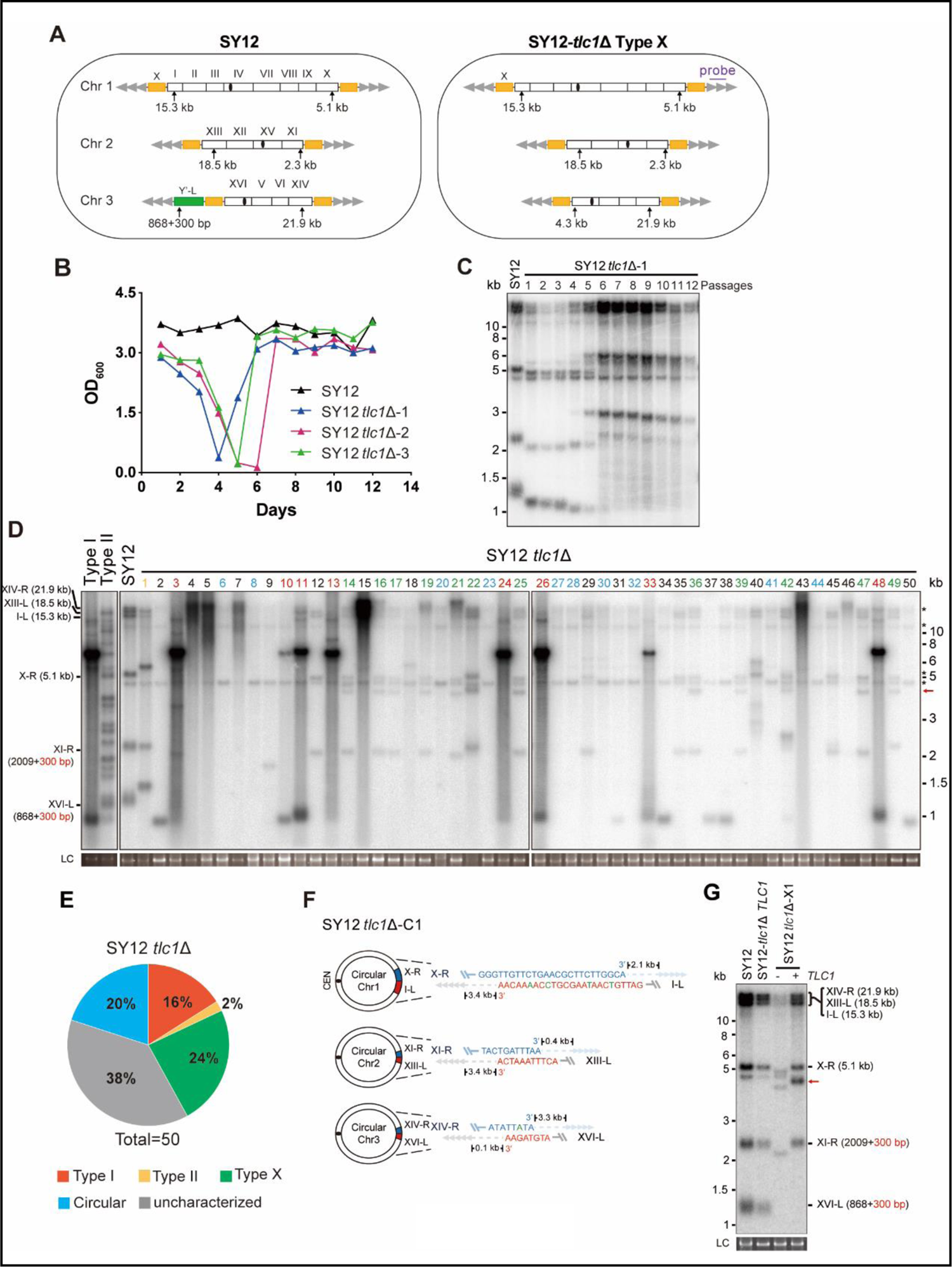
Survivor formation in SY12 *tlc1*Δ strain. (**A**) Schematic representation of chromosome (and telomere) structures (not drawn to scale) in the SY12 strain (left panel) and the Type X survivor (right panel). The Roman numerals, native chromosomes; the Arabic numerals on the left, chromosome numbers of SY12; yellow box, X-element; green box, Y’-element; tandem grey triangles, telomeres; black circles, centromere; vertical arrows and numbers, positions and lengths of the terminal Xhol digestion fragments detected by the telomeric TG1-3 probe. Chromosome numbers are omitted in the Type X survivor (right panel). (**B**) Cell viability assay in liquid medium. The growth of SY12 (labeled in black) and SY12 *tlc1*Δ (three clones labeled in blue, purple and green respectively) strains were monitored every 24 hr for 12 days. (**C**) Telomeric Southern blotting assay of SY12 *tlc1*Δ survivors. Genomic DNAs prepared from SY12 *tlc1*Δ survivors assayed in (B) were digested with XhoI and subjected to Southern blotting with a TG1-3 probe. (**D**) Telomere Southern blotting assay of SY12 *tlc1*Δ survivors obtained on solid medium. Genomic DNA from fifty independent SY12 *tlc1*Δ clones (labeled on top) was digested with XhoI and hybridized to a telomere-specific TG1–3 probe. Type II survivors: in orange; Type I survivors: in red; circular survivors: in blue; Type X survivors: in green; uncharacterized survivors: in black. Theoretical telomere restriction fragments of the SY12 strain are indicated on the left. The red arrows indicate the new band of about 4.3 kb emerged in Type X survivors. The asterisks indicate the non-specific bands. Genomic DNA stained with Gelred was used as a relative loading control (LC). (**E**) The ratio of survivor types in SY12 *tlc1*Δ strain. n=50; Type I, in red; Type II, in orange; Type X, in green; uncharacterized survivor, in grey; circular survivor, in blue. (**F**) Schematic of three circular chromosomes and fusion sequences in the SY12 *tlc1*Δ-C1 survivor. The sequence in blue indicates the sequences of X-R, XI-R or XIV-R, the sequence in red indicates the sequences of I-L, XIII-L or XVI-L. Bases in green are mis-paired. The numbers above or below the schematic line (chromosome) indicate the distance to the corresponding telomeres. (**G**) Telomere Southern blotting analysis of an SY12 *tlc1*Δ Type X survivor at the 20^th^ re-streak after *TLC1* reintroduction. The red arrows indicate the new band of about 4.3 kb emerged in Type X survivors. LC: loading control.

### Deletion of all of the X- and Y’-elements in the SY12 strain

We aimed to determine whether the subtelomeric X-elements are dispensable or not. In the SY12 strain, there are six X-elements distributed among six telomeres (Figure 2A, left panel). To precisely delete all X-and Y’-elements in SY12 strains, we employed a method that combines the efficient CRISPR-Cas9 cleavage system with the robust homologous recombination activity of yeast, as previously described (Shao et al., 2018; Shao et al., 2019). Briefly, the Cas9 nuclease cleaved the unique DNA sequences adjacent to the subtelomeric region (site S1) with the guidance of gRNA1. The resulting chromosome break was repaired through homologous recombination (HR) using the provided chromosome ends which excluding the X- and Y’-elements. Subsequently, the *URA3* marker and the guide RNA expression plasmid (pgRNA) were eliminated by inducing gRNA2 expression on pCas9 using galactose (Figure 3—figure supplement 1). This approach allowed us to initially delete the Y’-element and X-element in XVI-L, generating the SY12^YΔ^ strain (Figure 3A and supplementary file 4). Subsequently, through five successive rounds of deletions, we removed all remaining X-elements, resulting in the SY12^XYΔ^ strain (Figure 3A and supplementary file 4). To confirm the series of deletions, we performed PCR analysis using a primer located within the deletion region and another primer annealing upstream of the region (indicated by purple arrows in Figure 3— figure supplement 1). This analysis verified the complete deletion of the subtelomeric X- and Y’-elements (Figure 3B, rows 3 to 7). Additionally, we conducted a separate PCR analysis using primers specific to either X- or Y’-elements, which confirmed the absence of both X- and Y’-elements in the SY12^XYΔ^ strain (Figure 3B, rows 8). Subsequently, we inserted a Y’-long element (cloned from the native XVI-L sequence, which does not contain the centromere-proximal short telomere sequence) into the left arm of chromosome 3 in the SY12^XYΔ^ strain, resulting in the SY12^XYΔ+Y^ strain containing a single Y’-element but no X-element (Figure 3A and supplementary file 4). The successful insertion was confirmed by PCR analysis (Figure 3B, lane 9).

**Figure 3.**
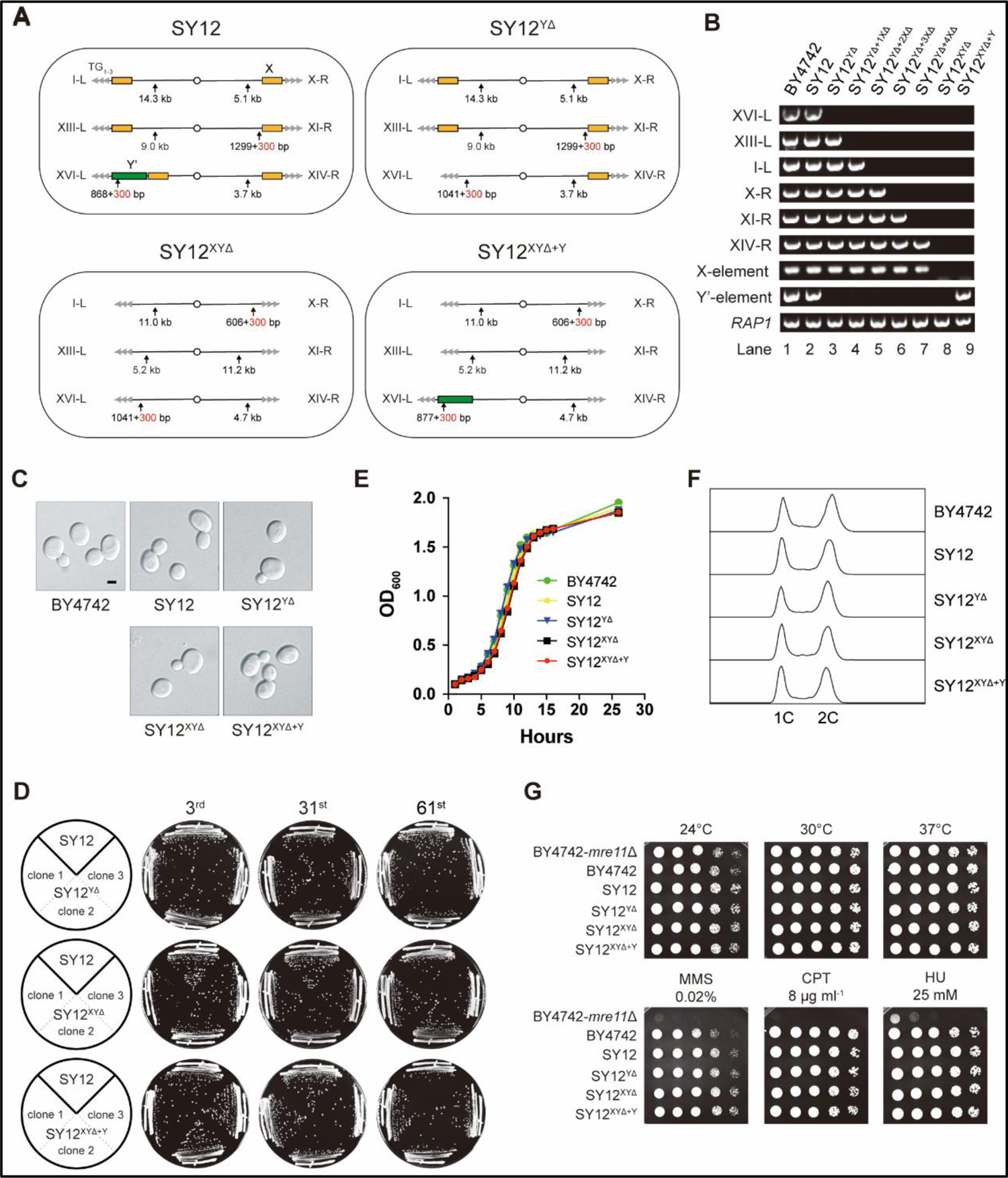
Characterization of SY12^YΔ^, SY12^XYΔ^ and SY12^XYΔ+Υ^ strains. (**A**) Schematic of chromosome structures in the SY12, SY12^YΔ^, SY12^XYΔ^ and SY12^XYΔ+Υ^ strains. Yellow box, X-element; green box, Y’-element; tandem grey triangles, telomeres. Vertical arrows and numbers indicate the positions and sizes of the sites and length of Xhol and PaeI-digested terminal fragments. (**B**) PCR analyses of the engineered sites of the individual telomeres (labeled on left) in BY4742, SY12, SY12^YΔ^, SY12^YΔ+1XΔ^, SY12^YΔ+2XΔ^, SY12^YΔ+3XΔ^, SY12^YΔ+4XΔ^, SY12^XYΔ^ and SY12^XYΔ+Υ^ strains (labeled on top). Primer sequences for the PCR analyses are listed in supplementary file 1. *RAP1* was an internal control. (**C**) Morphology of BY4742, SY12, SY12^YΔ^, SY12^XYΔ^ and SY12^XYΔ+Υ^ cells in the exponential growth phase (30°C in YPD). Shown are DIC images. Scale bar, 2 μm. (**D**) Growth analysis of the SY12, SY12^YΔ^, SY12^XYΔ^ and SY12^XYΔ+Υ^ strains. Several clones of the SY12, SY12^YΔ^, SY12^XYΔ^ and SY12^XYΔ+Υ^ strains were re-streaked on YPD plates 61 times at intervals of two days. Shown were the 3^rd^, 31^st^ and 61^st^ re-streaks. (**E**) Growth analysis of BY4742, SY12, SY12^YΔ^, SY12^XYΔ^ and SY12^XYΔ+Υ^ cells in liquid culture. Error bars represent standard deviation (s.d.), n = 3. (**F)** FACS analysis of DNA content of BY4742, SY12, SY12^YΔ^, SY12^XYΔ^ and SY12^XYΔ+Υ^ cells. (**G**) Dotting assays on YPD plates at low (24°C) and high (37°C) temperatures, or on YPD plates containing methyl methane sulfonate (MMS), camptothecin (CPT) or hydroxyurea (HU) at the indicated concentrations. The BY4742 *mre11*Δ haploid strain serves as a negative control because Mre11 is involved in the repair of double-stranded breaks (Lewis et al., 2004).

### Subtelomeric X- and Y’-elements are dispensable for cell proliferation, various stress responses, telomere length control and telomere silencing

The SY12^YΔ^, SY12^XYΔ^, and SY12^XYΔ+Y^ cells, cultured in YPD medium at 30°C, exhibited the same cell morphology as the parental strains SY12 and BY4742 (Figure 3C). To assess the stability of their genomes, we restreaked several clones of SY12^YΔ^, SY12^XYΔ^, and SY12^XYΔ+Y^ strains on YPD plates for a total of 61 times at two-day intervals (Figure 3D). Similar to the SY12 strain, the progeny colonies of SY12^YΔ^, SY12^XYΔ^, and SY12^XYΔ+Y^ grew robustly on solid medium (Figure 3D). Moreover, SY12^YΔ^, SY12^XYΔ^, and SY12^XYΔ+Y^ cells exhibited growth rates comparable to those of SY12 and BY4742 cells in liquid medium (Figure 3E). FACS analysis revealed that SY12^YΔ^, SY12^XYΔ^, and SY12^XYΔ+Y^ had the same 1C and 2C DNA content as wild-type cells (Figure 3F), indicating that the X- and Y’-elements are not necessary for cell proliferation under normal conditions. Additionally, the growth of SY12^YΔ^, SY12^XYΔ^, and SY12^XYΔ+Y^ cells at different temperatures (24 and 37°C) (Figure 3G, upper panel) closely resembled that of SY12 and BY4742 cells. Furthermore, SY12^YΔ^, SY12^XYΔ^, SY12^XYΔ+Y^, SY12, and BY4742 cells exhibited similar sensitivities to various genotoxic agents, including hydroxyurea (HU), camptothecin (CPT), and methyl methanesulfonate (MMS) (Figure 3G, lower panel). These results indicate that the X- and Y’-elements are dispensable for cellular responses to cold or heat treatment and DNA damage challenges, consistent with a recent study of “synthetic yeast genome project”, namely Sc2.0, showing that thousands of genome-wide edits, including the deletion of subtelomeric repetitive sequences, deletion of introns, and relocation of tRNAs genes, yielded a strain that displays comparable growth with wild type strain (Richardson et al., 2017; Zhao et al., 2023).

Next, we examined the effects of X- and Y’-element elimination on telomeres. Southern blotting assay revealed that SY12^YΔ^, SY12^XYΔ^, and SY12^XYΔ+Y^ cells maintained stable telomeres at a length of approximately 300 bp, comparable to that in SY12 cells (Figure 4A), indicating that the X- and Y’-elements are not required for telomere length regulation. To determine whether the deletion of X- and Y’-elements abolishes telomere silencing, we constructed haploid strains of SY12^YΔ^ *sir2*Δ, SY12^XYΔ^ *sir2*Δ, SY12^XYΔ+Y^ *sir2*Δ, SY12 *sir2*Δ, and BY4742 *sir2*Δ. We then performed real-time RT-PCR to quantify the expression of the *MPH3* and *HSP32* genes, located near the subtelomeric region of X-R (X-only end) and XVI-L (X-Y’ end), respectively (Figure 4B), and found that the increase of the *MPH3* or *HSP32* expression upon *SIR2* deletion in SY12^YΔ^, SY12^XYΔ^ and SY12^XYΔ+Y^ strains was more significant than that in the BY4742 or the SY12 strain, indicating that telomere silencing remains effective in the absence of X-and Y’-elements (Figure 4B). These findings align with previous studies showing that telomeres without an X- or Y’-element exert a position effect on the transcription of neighboring genes (Aparicio et al., 1991), and that X- and Y’-elements function as modulators of TPE (Fourel et al., 1999; Lebrun et al., 2001; Ottaviani et al., 2008).

In conclusion, the SY12^YΔ^, SY12^XYΔ^, and SY12^XYΔ+Y^ strains behave similarly to the wild-type SY12 strain under all tested conditions (Figures 3 and 4). Their simplified telomere structure makes them potentially useful tools for telomere studies.

**Figure 4.**
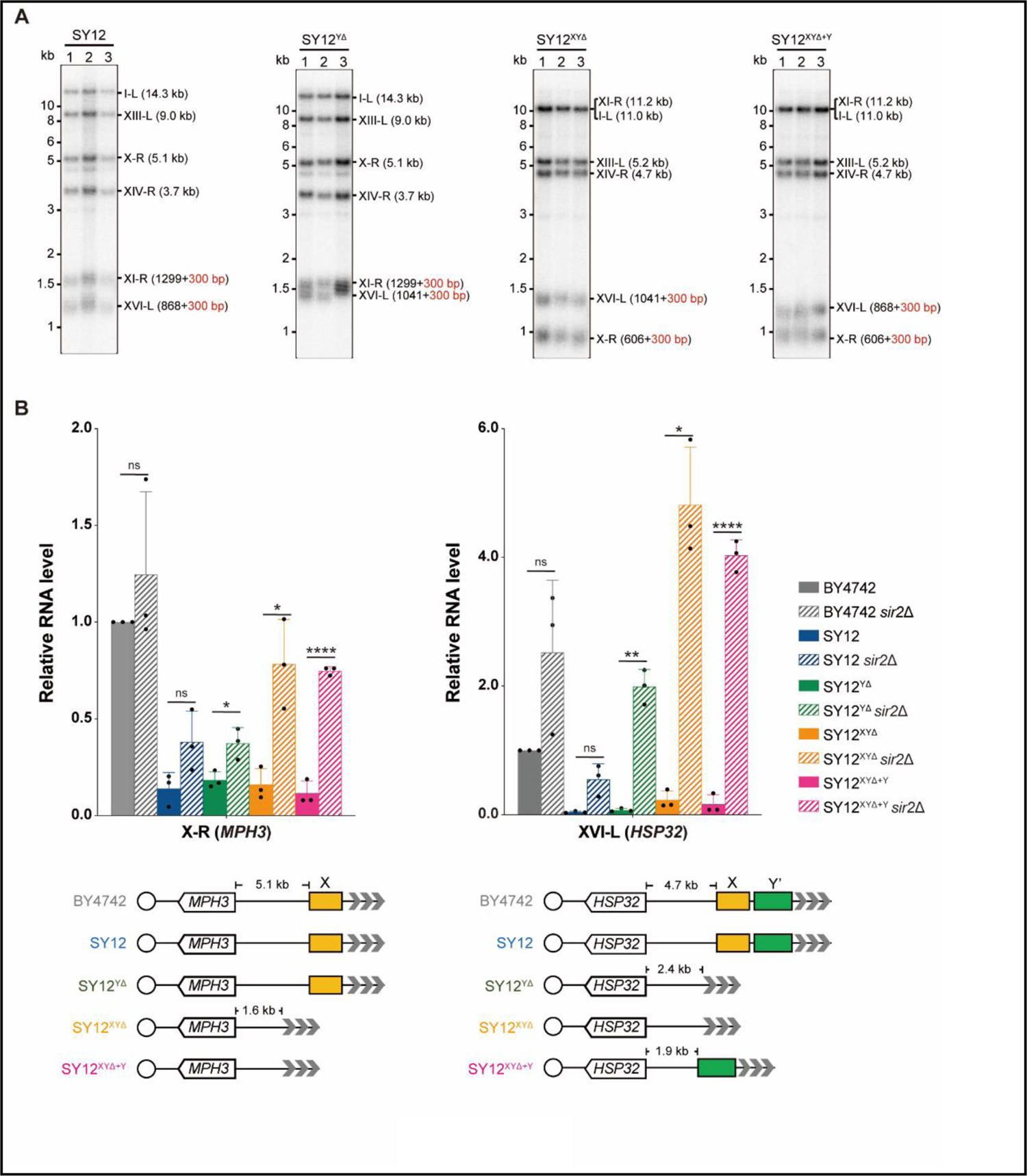
Telomere length and telomere silencing analyses of SY12^YΔ^, SY12^ΧΥΔ^ and SY12^XYΔ+Υ^ strains. (**A**) Southern blotting analysis of telomere length in SY12, SY12^YΔ^, SY12^XYΔ^ and SY12^XYΔ+Υ^ (labeled on top) cells. Genomic DNA prepared from three independent clones of SY12, SY12^YΔ^, SY12^XYΔ^ and SY12^XYΔ+Υ^ strains were digested with XhoI and PaeI, and then subjected to Southern blotting with a TG1-3 probe. The numbers in brackets indicate the telomere length of the corresponding chromosomes. (**B**) Expressions of *MPH3* and *HSP32* in ΒΥ4742, SY12, SY12^YΔ^, SY12^XYΔ^ and SY12^XYΔ+Υ^ cells were detected by qRT-PCR. The numbers above the schematic line (lower panels) indicate the distance to the corresponding subtelomeric elements or telomeres. The RNA levels of *MPH3* and *HSP32* were normalized by *ACT1*. The wild-type value is arbitrarily set to 1. Error bars represent standard deviation (s.d.), n = 3. ‘ns’, p> 0.5 (Student’s t-test); *, P < 0.05 (Student’s t-test); **, P < 0.01 (Student’s t-test).; ****, P < 0.0001 (Student’s t-test).

### Y’-elements are not strictly required for the formation of Type II survivors

The BY4742 strain harbors nineteen Y’-elements distributed among seventeen telomere loci. Numerous studies have emphasized the significance of Y’-elements in telomere recombination. For instance, Type I survivors exhibit significant amplification of Y’-elements (Lundblad and Blackburn, 1993; Teng and Zakian, 1999) and survivors show a marked induction of the potential DNA helicase Y’-Help1 encoded by Y’-elements (Yamada et al., 1998). Additionally, the acquisition of Y’-elements by short telomeres delays the onset of senescence (Churikov et al., 2014).

To investigate the requirement of Y’-elements in survivor formation, we deleted *TLC1* in SY12^YΔ^ cells and conducted a cell viability assay. The results demonstrated that three individual colonies underwent senescence and subsequently recovered at different passages in liquid media (Figure 5A). Further analysis through Southern blotting revealed that the telomeres of SY12^YΔ^ *tlc1*Δ cells underwent progressive shortening with each passage until reaching critically short lengths. Subsequently, TG_1-3_ recombination occurred, leading to abrupt telomere elongation (Figure 5B).

**Figure 5.**
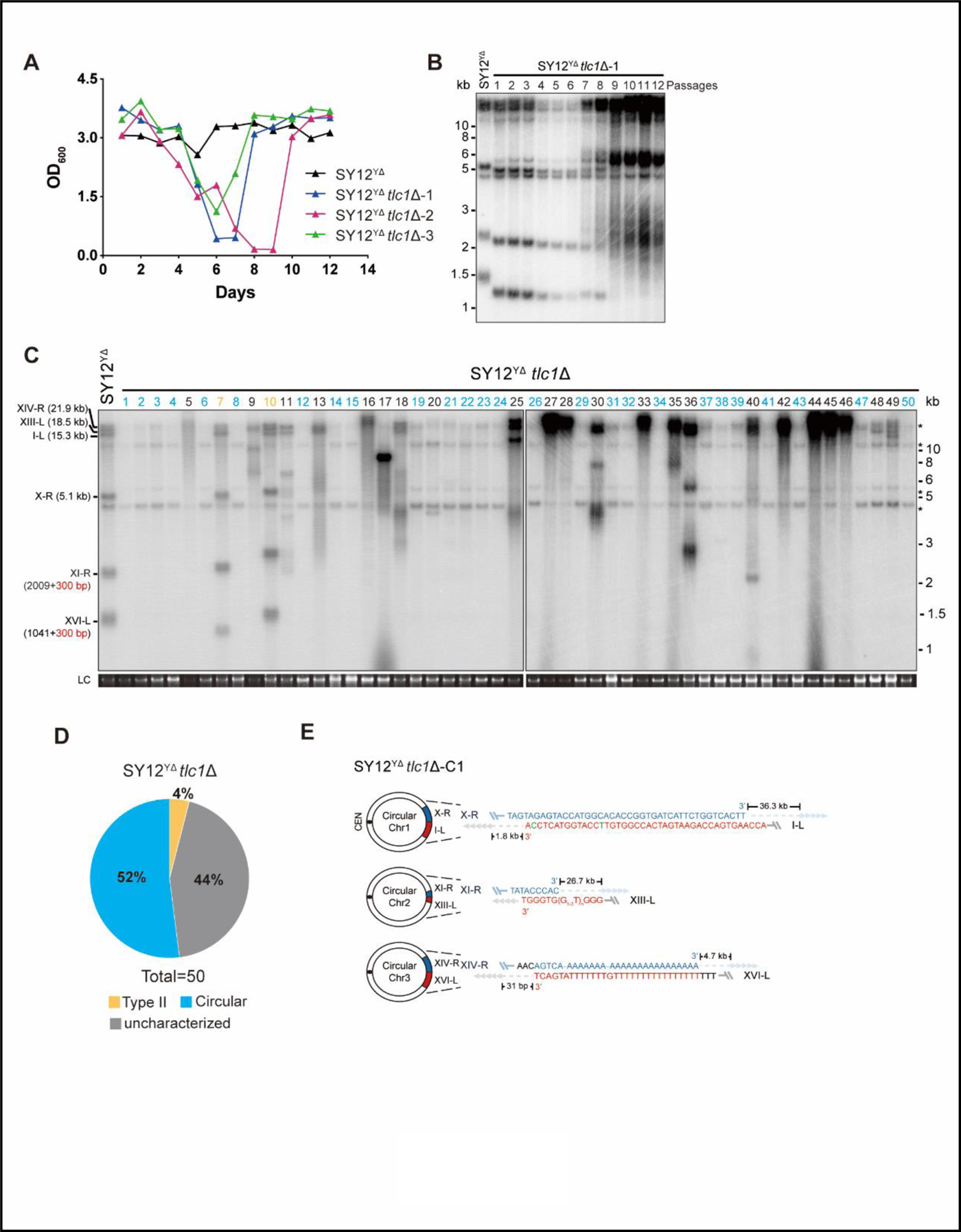
Survivor analysis of SY12^YΔ^ *tlc1*Δ strain. (**A**) Cell viability assay in liquid medium. The growth of SY12^YΔ^ (labeled in black) and SY12^YΔ^ *tlc1*Δ (three clones labeled in blue, purple and green respectively) strains were monitored every 24 hr for 12 days. (**B**) Telomeric Southern blotting assay of SY12^YΔ^ *tlc1*Δ survivors. Genomic DNAs prepared from SY12^YΔ^ *tlc1*Δ survivors assayed in (A) were digested with XhoI and subjected to Southern blotting with a TG1-3 probe. (**C**) Telomere Southern blotting analysis of SY12^YΔ^ *tlc1*Δ survivors obtained on solid medium. Genomic DNAs of fifty independent survivors (labeled 1 to 50 on top) were digested with XhoI and hybridized by a TG1-3 probe. Type II survivors: in orange; circular survivors: in blue; uncharacterized survivors: in black. Theoretical telomere restriction fragments of the SY12^YΔ^ strain are indicated on left. LC: loading control. (**D**) The ratio of survivor types in SY12 ^YΔ^ *tlc1*Δ strain. n=50; Type II, in orange; uncharacterized survivor, in grey; circular survivor, in blue. (**E**) Schematic of three circular chromosomes and fusion sequences in the SY12^YΔ^ *tlc1*Δ-C1 survivor. The sequence in blue indicates the sequences of X-R, XI-R or XIV-R, the sequence in red indicates the sequences of I-L, XIII-L or XVI-L. Bases in green are mis-paired, dashes are deleted. The numbers above or below the schematic line (chromosome) indicate the distance to the corresponding telomeres.

Next, we examined the telomere patterns of fifty independent SY12^YΔ^ *tlc1*Δ survivors using a multiple-colony streaking assay and Southern blotting analysis. Out of the fifty clones analyzed, no Type I survivors were detected due to the deletion of Y’-elements in SY12^YΔ^ strain (Figure 5C). Two clones (labeled in orange, 4% of the survivors tested) displayed heterogeneous telomere tracts (Figure 5C and Figure 5D). Reintroduction of *TLC1* into a representative clone (named SY12^YΔ^ *tlc1*Δ-T1) resulted in telomere length restoration similar to SY12^YΔ^ cells (Figure 5—figure supplement 1A), indicating their classification as Type II survivors. Twenty-six clones (labeled in blue, 52% of the survivors tested) exhibited patterns identical to that of the SY12 *tlc1*Δ circular survivors (Figure 5C, Figure 5D and Figure 2D). Further mapping of erosion borders and sequencing of fusion junctions (Figure 5E, Figure 5—figure supplement 2, and supplementary file 3) confirmed that three chromosomes from a randomly selected clone (named SY12^YΔ^ *tlc1*Δ-C1) underwent intra-chromosomal fusions mediated by microhomology sequences. The erosion sites and fusion sequences differed from those observed in SY12 *tlc1*Δ-C1 cells (Figure 2F), suggesting the stochastic nature of intra-chromosome end fusion by MMEJ. As expected, the telomere Southern blotting pattern (XhoI digestion) of the SY12^YΔ^

*tlc1*Δ-C1 survivor remained unchanged following telomerase reintroduction (Figure 5—figure supplement 1B). Further PFGE analysis confirmed that the chromosomes in SY12^YΔ^ *tlc1*Δ-C1 were circulated (Figure 2—figure supplement 5). Notably, a significant proportion of the survivors displayed telomere signals that were different from those of either the Type II or circular survivors (labeled in black, 44% of the survivors tested, Figure 5C and Figure 5D), and they were uncharacterized survivors. Further deletion of *RAD52* in the SY12^YΔ^ *tlc1*Δ cells affected, but did not eliminate, survivor generation (Figure 5—figure supplement 3A). Southern blotting assay confirmed that most of the recovered clones were circular survivors, and two were uncharacterized survivors (clone 9 and 16, labeled in black, Figure 5—figure supplement 3B) suggesting that survivor formation in SY12^YΔ^ *tlc1*Δ rad52Δ cells does not strictly rely on the homologous recombination. Overall, these findings indicate that Y’-elements are not strictly required for Type II survivor formation (Churikov et al., 2014).

### X-elements are not strictly necessary for survivor generation

To investigate the contribution of X-elements to telomere recombination, we employed the SY12^XYΔ+Y^ strain, which contains only one Y’-element in the subtelomeric region, and the SY12^XYΔ^ *tlc1*Δ strain, which lacks both the X- and Y’-elements. Subsequently, we deleted *TLC1* in the SY12^XYΔ+Y^ and SY12^XYΔ^ strains and conducted a cell viability assay. Consistently, the deletion of *TLC1* in SY12^XYΔ+Y^ and SY12^XYΔ^ resulted in telomere shortening, senescence, and the formation of Type II survivors (Figure 6—figure supplement 1). Then, fifty independent clones of SY12^XYΔ+Y^ *tlc1*Δ or SY12^XYΔ^ *tlc1*Δ survivors were examined using Southern blotting (Figure 6A and 6B).

Among the SY12^XYΔ+Y^ survivors analyzed, twenty-two clones underwent chromosomal circularization (labeled in blue, 44% of the survivors tested, Figure 6A and Figure 6C). We randomly selected a clone named SY12^XYΔ+Y^ *tlc1*Δ-C1, and the results of erosion-border mapping and fusion junction sequencing showed that it had undergone intra-chromosomal fusions mediated by microhomology sequences (Figure 6D, Figure 6—figure supplement 2, and supplementary file 3). Subsequently, Southern blotting revealed that the chromosome structure of SY12^XYΔ+Y^ *tlc1*Δ-C1 remained unchanged after *TLC1* reintroduction (Figure 6— figure supplement 3), and PFGE analysis confirmed the circular chromosome structure in SY12^XYΔ+Y^ *tlc1*Δ-C1 (Figure 2—figure supplement 5). Additionally, seven clones utilized the Type II recombination pathway and exhibited heterogeneous telomeric TG_1-3_ tracts (labeled in orange, 14% of the survivors tested, Figure 6A and Figure 6C). Reintroduction of *TLC1* into a representative clone (named SY12^XYΔ+Y^ *tlc1*Δ-T1) restored the telomere length to normal (Figure 6—figure supplement 3). These findings indicate that the majority of cells underwent intra-chromosomal circularization or TG_1-3_ recombination. While even though there is a Y’-element, no Type I survivors were generated in SY12^XYΔ+Y^ *tlc1*Δ survivors (Figure 6A). We speculated that the short TG_1-3_ repeats located centromere-proximal to the Y’-elements play a crucial role in strand invasion and subsequent Y’-recombination. This speculation is consistent with a previous report stating that Type I events are virtually absent in the yeast strain Y55, which lacks TG_1-3_ repeats centromere-proximal to the Y’-element (Louis, 2001). We also observed some clones displayed non-canonical telomere signals like SY12 *tlc1*Δ “uncharacterized” survivors (labeled in black, 42% of the survivors tested, Figure 6A and Figure 6C). Overall, these data suggest that X-elements are not strictly necessary for survivor formation.

Among the SY12^XYΔ^ survivors, twenty-four displayed a “circular survivor” pattern (labeled in blue, 48% of the survivors tested, Figure 6B and Figure 6E). Additional PCR-sequencing assays and PFGE analysis of the SY12^XYΔ^ *tlc1*Δ-C1 cells confirmed the occurrence of intra-chromosomal fusions mediated by microhomology sequences (Figure 6F, Figure 6—figure supplement 4, supplementary file 3 and Figure 2—figure supplement 4). Reintroduction of *TLC1* into a representative clone named SY12^XYΔ^ *tlc1*Δ-C1 could restore its telomere length to WT level (Figure 6—figure supplement 5A). Four of fifty survivors harbored Type II telomere structure (labeled in orange, 8% of the survivors tested, Figure 6B and Figure 6E). Reintroduction of *TLC1* into a representative clone named SY12^XYΔ^ *tlc1*Δ-T1 could restore its telomere length to WT level (Figure 6—figure supplement 5B). Some of the survivors (labeled in black, 44% of the survivors tested, Figure 6B and Figure 6E) were not characterized. Like in SY12 *tlc1*Δ cells, Rad52 is not strictly required for the formation of circular survivors in SY12^XYΔ^ *tlc1*Δ *rad52*Δ and SY12^XYΔ+Y^ *tlc1*Δ *rad52*Δ strains (Figure 6—figure supplement 6A and B). To investigate whether Type I-specific mechanisms are still utilized in the survivor formation in Y’-less strain, we deleted *RAD51* in SY12^XYΔ^ *tlc1*Δ, and found that SY12^XYΔ^ *tlc1*Δ *rad51*Δ strain was able to generate three types of survivors, including Type II survivor, circular survivor and uncharacterized survivor (Figure 6—figure supplement 7A), similar to the observations in SY12^XYΔ^ *tlc1*Δ strain (Figure 6B). Notably, the proportions of circular and uncharacterized survivors in the SY12^XYΔ^ *tlc1*Δ *rad51*Δ strain were 36% (9/25) and 32% (8/25) (Figure 6—figure supplement 7B and supplementary file 5), respectively, lower than 48% and 44% in the SY12^XYΔ^ *tlc1*Δ strain (Figure 6E and supplementary file 5). Accordingly, the ratio of Type II survivor in SY12^XYΔ^ *tlc1*Δ *rad51*Δ was (32% of the survivors tested, Figure 6—figure supplement 7B and supplementary file 5) was higher than SY12^XYΔ^ *tlc1*Δ strain (8% of the survivors tested, Figure 6E and supplementary file 5), suggesting that Type I-specific mechanisms still contribute to the survivor formation even in the Y’-less strain SY12^XYΔ^. Collectively, the aforementioned data suggest that X-elements, as well as Y’-elements, are not essential for the generation of Type II survivors.

**Figure 6.**
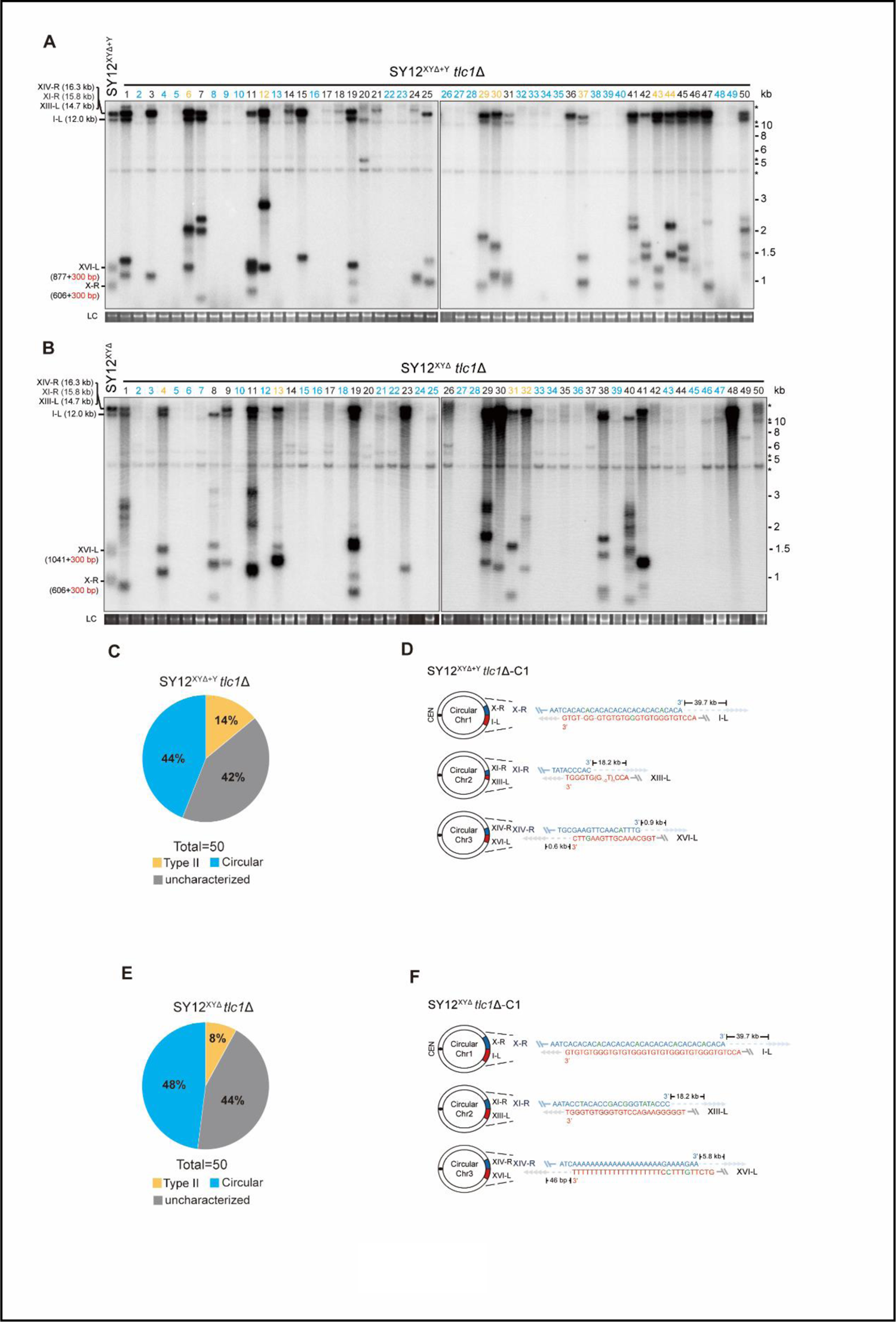
Survivor analysis of SY12^XYΔ^ *tlc1*Δ and SY12^XYΔ+Y^ *tlc1*Δ strains. (**A**) and (**B**) Telomere Southern blotting analysis of SY12^XYΔ+Y^ *tlc1*Δ (A) and SY12^XYΔ^ *tlc1*Δ (B) survivors obtained on solid medium. fifty independent survivors (labeled 1 to 50 on top) were randomly picked, and their genomic DNAs were digested with XhoI and subjected to the Southern blotting assay with a TG1-3 probe. Type II survivors: in orange; circular survivors: in blue; uncharacterized survivors: in black. The sizes of individual telomere restriction fragments of the SY12^XYΔ+Y^ and SY12^XYΔ^ strain are indicated on the left. LC: loading control. (**C**) and (**E**) The percentage of survivor types in SY12 ^XYΔ+Y^ *tlc1*Δ (C) and SY12 ^XYΔ^ *tlc1*Δ (E) strains. n=50; Type II, in orange; uncharacterized survivor, in grey; circular survivor, in blue. (**D**) and (**F**) Schematic of three circular chromosomes and fusion sequences in the SY12^XYΔ+Y^ *tlc1*Δ-C1 (D) and SY12^XYΔ^ *tlc1*Δ-C1 (F) survivors, respectively. The sequence in blue indicates the sequences of X-R, XI-R or XIV-R, the sequence in red indicates the sequences of I-L, XIII-L or XVI-L. Bases in green are mis-paired, dashes are deleted. The numbers above or below the schematic line (chromosome) indicate the distance to the corresponding telomeres.

## Discussion

The wild-type yeast strain BY4742, commonly used in laboratories, possesses nineteen Y’-elements at seventeen telomere loci and thirty-two X-elements at thirty-two telomere loci. This abundance of Y’-elements and X-elements poses challenges for loss-of-function studies, highlighting the need for a strain lacking all Y’-elements and X-elements. Fortunately, we have previously constructed the single-chromosome yeast strain SY14, which contains only one copy of Y’-element and two copies of X-element (Shao et al., 2018), and could have been an ideal tool. However, the telomerase-null survivors of SY14 mainly bypassed senescence through chromosomal circularization, providing limited insights into the roles of Y’-and X-elements in telomere maintenance (Wu et al., 2020). Therefore, in this study, we employed the SY12 strain, which has three chromosomes, to investigate the functions of Y’- and X-elements at telomeres (Figure 2A, left panel).

We constructed the SY12^YΔ^, SY12^XYΔ+Y^, and SY12^XYΔ^ strains, which lack the Y’-element, X-elements, and both X- and Y’-elements, respectively (Figure 3A). Surprisingly, the SY12^YΔ^, SY12^XYΔ^, and SY12^XYΔ+Y^ strains exhibited minimal defects in cell proliferation, genotoxic sensitivity, and telomere homeostasis (Figures 3 and 4). These results demonstrate, for the first time, that both X- and Y’-elements are dispensable for cellular functions. Thus, the SY12^YΔ^, SY12^XYΔ^, and SY12^XYΔ+Y^ strains established in this study, with their simplified telomere structures, are valuable resources for telomere biology research.

Subtelomeric regions are known to be highly variable and often contain species-specific homologous DNA sequences. In the case of fission yeast, subtelomeric regions consist of subtelomeric homologous (SH) and telomere-distal sequences. Previous studies have shown that subtelomeric homologous sequences in fission yeast do not significantly impact telomere length, mitotic cell growth, or stress responses. However, they do play a role in buffering against the spreading of silencing signals from the telomere (Tashiro et al., 2017). Though the “core X” sequence acts as a protosilencer (Lebrun et al., 2001), the X-STRs and Y’-STAR possess anti-silencing properties that limit the spreading of heterochromatin in budding yeast (Fourel et al., 1999), the telomere position effect remains effective in the strains that lack both X- and Y’-elements (Figure 4B). Given the remarkable differences in both sequence and size between the subtelomeric regions of budding yeast and fission yeast, it is difficult to compare the extent to which subtelomeric elements affect telomere silencing.

Amplification of Y’-element(s) is a characteristic feature of canonical Type I survivors. Type I survivor emerged in SY12 strain indicating that multiple Y’-elements in tandem is not strictly required for type I recombination (Figure 2D). Interestingly, the telomerase-null SY12^YΔ^ and SY12^XYΔ^ cells, lacking Y’-elements, failed to generate Type I survivors but could generate Type II survivors, indicating that the acquisition of Y’-elements is not a prerequisite for Type II survivor formation (Figure 5C and Figure 6B). These observations support the notion that Type I and Type II survivors form independently, although both may utilize a common alternative telomere-lengthening pathway (Kockler et al., 2021). Moreover, a subset of SY12 *tlc1*Δ, SY12^YΔ^ *tlc1*Δ, SY12^XYΔ+Y^ *tlc1*Δ, and SY12^XYΔ^ *tlc1*Δ cells could escape senescence and become survivors through microhomology-mediated intra-chromosomal end-to-end fusion (chromosome circularization) (Figure 2D, Figure 5C, Figure 6A and 6B, labeled in blue). Notably, the survivors with all circular chromosomes were readily recovered from the telomerase-null SY11 to SY14, but not SY1 to SY10 cells (Figure 1). Several reasons could account for this. First, a smaller number of telomeres provides fewer recombination donors and acceptors, resulting in less efficient inter-chromosomal homologous recombination (e.g., TG_1-3_ tracts recombination or Y’-element acquisition). Second, the continuously shortened telomeres of linear chromosomes may trigger another round of senescence, while survivors with circular chromosomes do not encounter end-replication problems and therefore exhibit greater stability. Third, the presence of homologous sequences at both chromosome ends appears to be a minimum requirement for microhomology-mediated intra-chromosomal end-to-end fusion. With fewer homologous sequences, the probability of chromosome circularization decreases, and with more chromosomes, the likelihood of circularizing each chromosome within a cell diminishes. Fourth, in cells with fewer telomeres, intra-chromosomal telomere fusions are more likely to occur, while lethal inter-chromosomal fusions are competed out. However, we can speculate that in telomerase-null cells with eroded chromosome ends, stochastic repair mechanisms such as homologous recombination, microhomology-mediated end joining, and inter- and intra-chromosomal fusions operate simultaneously. Only those survivors that maintain a relatively stable genome and robust growth can be experimentally recovered.

*Saccharomyces cerevisiae* (budding yeast) and *Schizosaccharomyces pombe* (fission yeast) are the most commonly used laboratory systems, separated by approximately 1 Gya (billion years ago) according to molecular-clock analyses (Hedges, 2002). Despite both species having genomes are both over 12 megabases in length, haploid *S. cerevisiae* contains 16 chromosomes, while *S. pombe* has only 3 chromosomes (Forsburg, 2005). The telomerase-independent mechanisms for maintaining chromosome ends differ between these two yeasts. In budding yeast, homologous recombination is the primary mode of survival in telomerase-deficient cells, resulting in the generation of Type I or Type II survivors (McEachern and Haber, 2006). Telomerase- and recombination-deficient cells occasionally escape senescence through the formation of palindromes at chromosome ends in the absence of *EXO1* (Maringele and Lydall, 2004). Fission yeast cells lacking telomerase can also maintain their chromosome termini by recombining persistent telomere sequences, and survivors with all intra-circular chromosomes (Nakamura et al., 1998) or intermolecular fusions (Tashiro et al., 2017; Wang and Baumann, 2008) have been observed. In our research, some SY12 *tlc1*Δ cells, which have three chromosomes, also bypassed senescence by circularizing their chromosomes (Figure 2D), suggesting that a lower chromosome number increases the likelihood of recovering survivors containing circular chromosomes.

While most eukaryotes employ telomerase for telomere replication, some eukaryotes lack telomerase and utilize recombination as an alternative means to maintain telomeres (Biessmann and Mason, 1997). In *Drosophila*, telomeres are replicated through a retrotransposon mechanism (Levis et al., 1993; Louis, 2002). The structure and distribution of Y’-elements in *S. cerevisiae* suggest their origin from a mobile element (Jager and Philippsen, 1989; Louis and Haber, 1992), and Y’-elements can be mobilized through a transposition-like RNA-mediated process (Maxwell et al., 2004). In telomerase-deficient yeast cells, homologous recombination can acts as a backup mechanism for telomere replication (Lundblad and Blackburn, 1993), and the reintroduction of telomerase efficiently inhibits telomere recombination and dominates telomere replication (Chen et al., 2009; Peng et al., 2015; Teng and Zakian, 1999), These findings suggest that subtelomeric region amplification mediated by recombination and/or transposition may represent ancient telomere maintenance mechanisms predating the evolution of telomerase (de Lange, 2004). Therefore, subtelomeric X- and Y’-elements might be considered as evolutionary “fossils” in the *S. cerevisiae* genome, and their elimination has little impact on telomere essential functions and genome stability.

## MATERIALS AND METHODS

### Yeast strains and plasmids

Yeast strains used in this study are listed in supplementary file 6. The plasmids for gene deletion and endogenous expression of *TLC1* were constructed based on the pRS series as described previously (Sikorski and Hieter, 1989). We use PCR to amplify the upstream and downstream sequence adjacent to the target gene, and then the PCR fragments were digested with different restriction enzymes and inserted into pRS plasmids. Plasmids were introduced into budding yeast by standard procedures, and transformants were selected on auxotrophic medium (Orr-Weaver et al., 1981).

### Multiple-colony streaking assay

Single clones of indicated yeast strains were randomly picked and streaked on extract-peptone-dextrose (YPD) plates. Thereafter, several clones of their descendants were passaged by successive re-streaks at 30°C. This procedure was repeated dozens of times every two days.

### Telomere Southern blotting

Southern blotting was performed as previously described (Hu et al., 2013). Yeast genomic DNA was extracted by a phenol chloroform method. Restriction fragments were separated by electrophoresis in 1% agarose gel, transferred to Amersham Hybond-N^+^ membrane (GE Healthcare) and hybridized with α-^32^P dCTP labeled probe.

### Cell viability assay

Cell viability assay was performed as previously described with few modifications (Le et al., 1999). Three independent single colonies of indicated strains were grown to saturation at 30°C. Then the cell density was measured every 24 hours by spectrometry (OD_600_), and the cultures were diluted to the density at OD_600_ = 0.01. This procedure was repeated several times to allow the appearance of survivors. The genomic DNA samples at indicated time points were harvested for telomere length analysis.

### Molecular analysis of circular chromosomes

Fusion events were determined by PCR amplification and DNA sequencing. Genomic DNA was extracted by phenol chloroform. Frist, we use primers pairs located at different sites of each chromosome arm at an interval of 1 kb (listed in Supplementary file 1) to determine the erosion site of each chromosome; PCR was performed as standard procedures in 10 μl reactions by TaKaRa Ex Taq. To amplify the sequence of fusion junction we use pairs of primers oriented to different arm of each chromosome; PCR was performed as standard procedures in 50 μl reactions by TaKaRa LA Taq. The fragments were purified by kit (QINGEN), then they were sequenced directly or cloned into the pMD18-T Vector (TaKaRa) for sequencing.

### CRISPR-Cas9 mediated X- and Y’-elements deletion

X- and Y’-elements were deleted as described (Shao et al., 2018; Shao et al., 2019). Briefly, pgRNA and a DNA targeting cassette, containing a selection marker, a homology arm (DR1), a direct repeat (DR2), and telomeric repeats, were co-introduced into indicated cells harboring pCas9. pCas9 nuclease was directed to a specific DNA sequence centromere-proximal to the subtelomeric region with the guidance of gRNA1, where it induces a double-stranded break. Homologous recombination between the broken chromosome and the provided DNA targeting cassette caused the deletion of X- and Y’-elements. The positive transformants identified by PCR were transferred into the galactose-containing liquid medium, which induces the expression of the gRNA2 on pCas9 to cut at the target site near the *URA3* gene and on the backbone of pgRNA. Then the culture was plated on the medium containing 5’-FOA to select for eviction of the *URA3* marker.

### Cell growth assay

Three individual colonies of the indicated strains were inoculated into 5 ml liquid medium and incubated at 30°C. The cell cultures were then diluted in 30 ml of fresh YPD medium to the density at OD_600_ = 0.1. Then the density of cells was measured by spectrometry (OD_600_) hourly.

### Fluorescence-activated cell sorting (FACS) assay

The FACS analysis was performed as previously described (He et al., 2019). Yeast cells were cultured at 30°C until the log phase, and then 1 ml of the cells was harvested. The cells were washed with cold sterile ddH_2_O and fixed with 70% ethanol overnight at 4°C. The following day, the cells were washed with 50 mM sodium citrate buffer (pH 7.2) and then digested with 0.25 mg/ml RNase A at 37°C for 2-3 hours, followed by 0.2 mg/ml Protease K at 50°C for 1 hour. Both RNase A and Protease K were diluted in sodium citrate buffer. The cells were resuspended in 500 μl sodium citrate buffer and then sonicated for 45 seconds at 100% power. The DNA of the cells was stained with 20 μg/ml propidium iodide (PI) at 4°C overnight or at room temperature for 1 hour. FACS analysis was performed on a BD LSRII instrument.

### Serial dilution assay

A single colony per strain was inoculated into 3 ml liquid medium and incubated at 30°C. The cell cultures were then adjusted to a concentration of OD_600_ ∼0.5. Five-fold serially diluted cells were spotted on the indicated plates. The plates were incubated at 30°C for the appropriate time prior to photography.

### RNA extraction and RT-qPCR

Three independent single colonies of indicated strains were grown to log phase at 30°C. Yeast pellets from a 1 ml cell culture were digested with Zymolyase 20 T (MP Biomedicals, LLC) to obtain spheroplasts. RNA was extracted with RNeasy mini kit (Qiagen) followed by reverse transcription using the Fastquant RT kit (Tiangen). Real-time PCR was carried out using SYBR Premix Ex Taq^TM^ II (Takara) on the Applied Biosystems StepOne Real-Time PCR System. Primer pairs used in RT-qPCR were listed in supplementary file 1. The gene expression levels were normalized to that of *ACT1* and the wild-type value is arbitrarily set to 1.

### Pulsed-field gel electrophoresis (PFGE) analysis

DNA plugs for pulse-field gel electrophoresis (PFGE) were prepared according to the manufacturer’s instructions (Bio-Rad) and Ishii et al (Ishii et al., 2008). Fresh yeast cells were inoculated in 50 ml YPD and incubated at 30°C until the OD600 reached approximately 1.0. The cells were subsequently harvested, washed twice with cold EDTA buffer (50 mM, pH 8.0), and resuspended in 300 μl of CSB buffer (10 mM pH 7.2 Tris-Cl, 20 mM NaCl, 100 mM pH 8.0 EDTA, 4 mg/ml lyticase) and blended with 300 μl of 2% low-melt agarose (Bio-Rad). Then, 100 μl of resuspended cells were added to each plug and incubated at 4°C for 30 min until the agarose plugs were solidified. The solidified agarose plugs were incubated in lyticase buffer (10 mM pH 7.2 Tris-Cl, 100 mM pH 8.0 EDTA, 1 mg/ml lyticase) at 37°C for 3 hours, followed by incubation in Proteinase K Reaction Buffer (100 mM pH 8.0 EDTA, 0.2% sodium deoxycholate, 1% sodium lauryl sarcosine) containing 1 mg/ml Proteinase K at 50°C for 12 hours. The plugs were washed four times in 25 ml of wash buffer (20 mM Tris, pH 8.0, 50 mM EDTA) for 1 hour each time at room temperature with gentle agitation. The plugs were then fixed into a pulsed field agarose gel (Bio-Rad), and the CHEF-DR II Pulsed Field

Electrophoresis System (Bio-Rad) was used for gel electrophoresis. The electrophoresis conditions for separation were as follows: 0.8% agarose gel, 1×TBE buffer, 14°C temperature, first run: initial switch time 1200 s; final switch time 1200 s; run time 24 hours; voltage gradient 2 V/cm; angle 96°; second run: initial switch time 1500 s; final switch time 1500 s; run time 24 hours; voltage gradient 2 V/cm; angle 100°; third run: initial switch time 1800 s; final switch time 1800 s; run time 24 hours; voltage gradient 2 V/cm; angle 106°. The gel was stained with GelstainRedTM nucleic acid dye (US EVERBRIGHT), and PFGE Gels were imaged by Tanon 2500.

## ACKNOWLEDGMENTS

We thank members of Zhou lab for discussions and suggestions for this project.

## AUTHOR CONTRIBUTIONS

Conceptualization: Zhi-Jing Wu, Jin-Qiu Zhou.

Funding acquisition: Jin-Qiu Zhou.

Investigation: Can Hu, Zhi-Jing Wu, Xue-Ting Zhu.

Methodology: Zhi-Jing Wu, Yangyang Shao, Zhongjun Qin.

Supervision: Jin-Qiu Zhou.

Writing-original draft: Zhi-Jing Wu.

Writing-review & editing: Zhi-Jing Wu, Can Hu, Xue-Ting Zhu, Ming-Hong He, Jin-Qiu Zhou.

## Figure supplements

**Figure 2—figure supplement 1.**
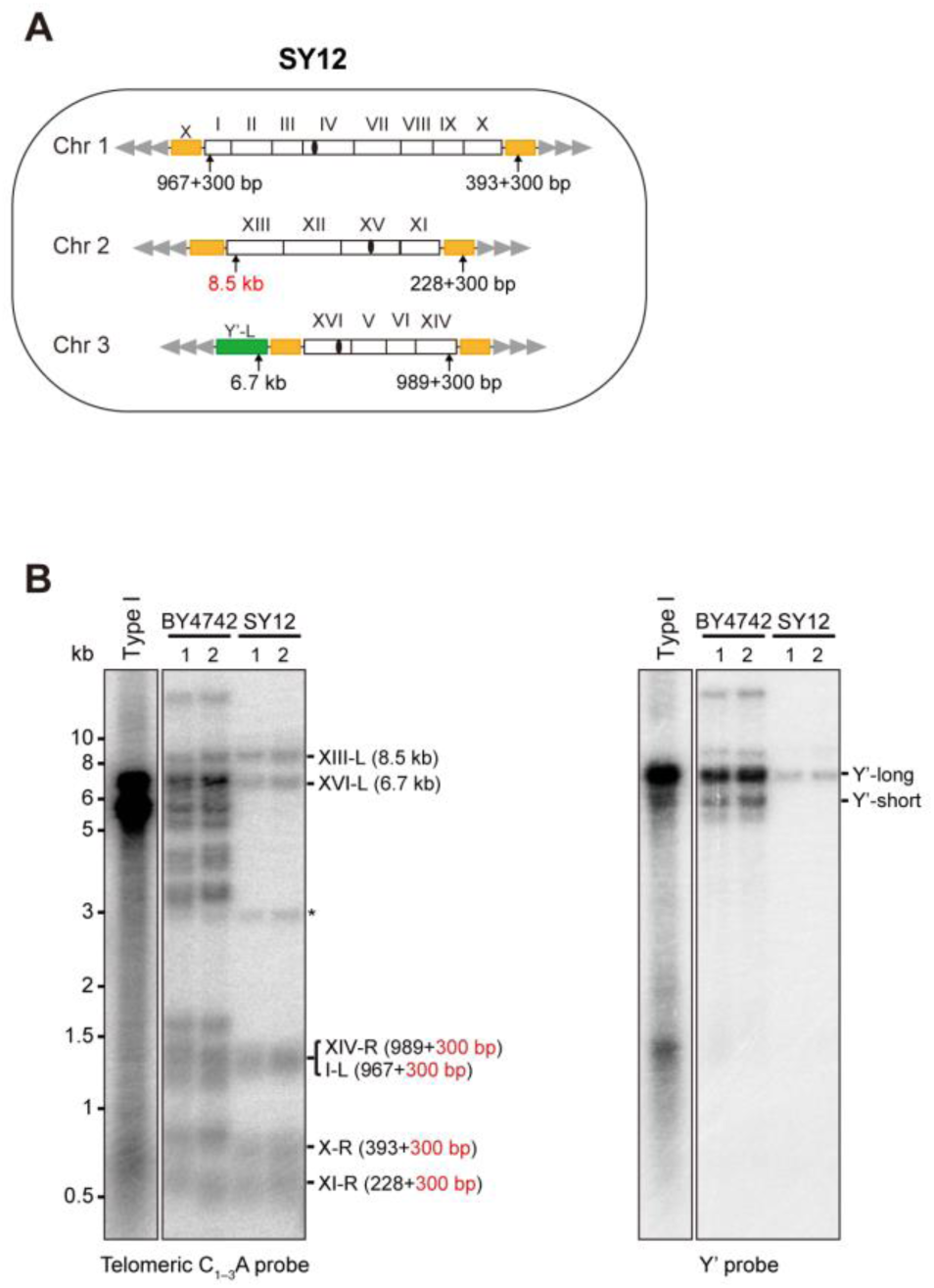
Characterization of SY12 strain. (**A**) Schematic representation of chromosome (and telomere) structures (not drawn to scale) in the SY12 strain. The Roman numerals, native chromosomes; the Arabic numerals on the left, chromosome numbers of SY12; yellow box, X-element; green box, Y’-element; tandem grey triangles, telomeres; black circles, centromere; vertical arrows and numbers, positions and lengths of the terminal Ndel digestion fragments detected by the telomeric TG1-3 probe. (**B**) Southern blotting analysis of telomere length in BY4742 and SY12 (labeled on top) cells. Genomic DNA prepared from two independent clones of BY4742 and SY12 strains was digested with NdeI, and then subjected to Southern blotting with a TG1-3 probe (left panel). The numbers in brackets indicate the telomere length of the corresponding chromosomes. The blot was then stripped and reprobed with a Y’ probe (right panel). The asterisk indicates the non-specific band.

**Figure 2—figure supplement 2.**
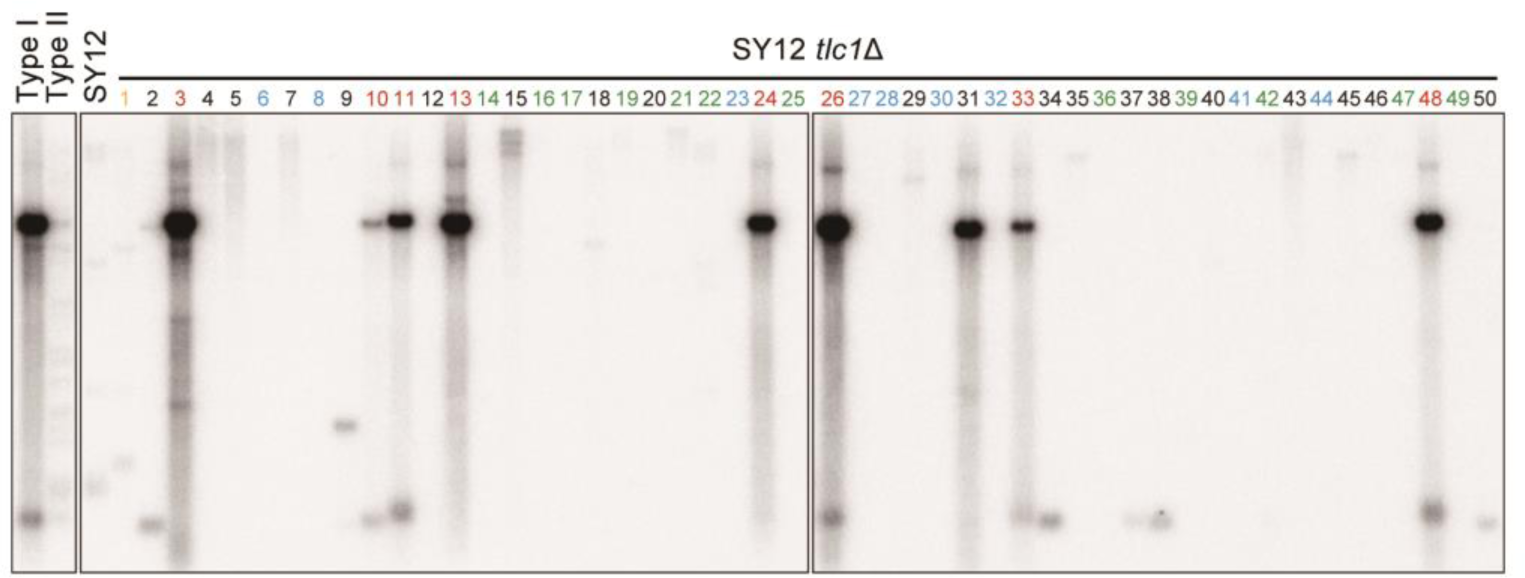
Telomere Southern blot with a Y’-element probe examining SY12 *tlc1*Δ survivors. The blot in Figure 2D was then stripped and reprobed with a Y’ probe.

**Figure 2—figure supplement 3.**
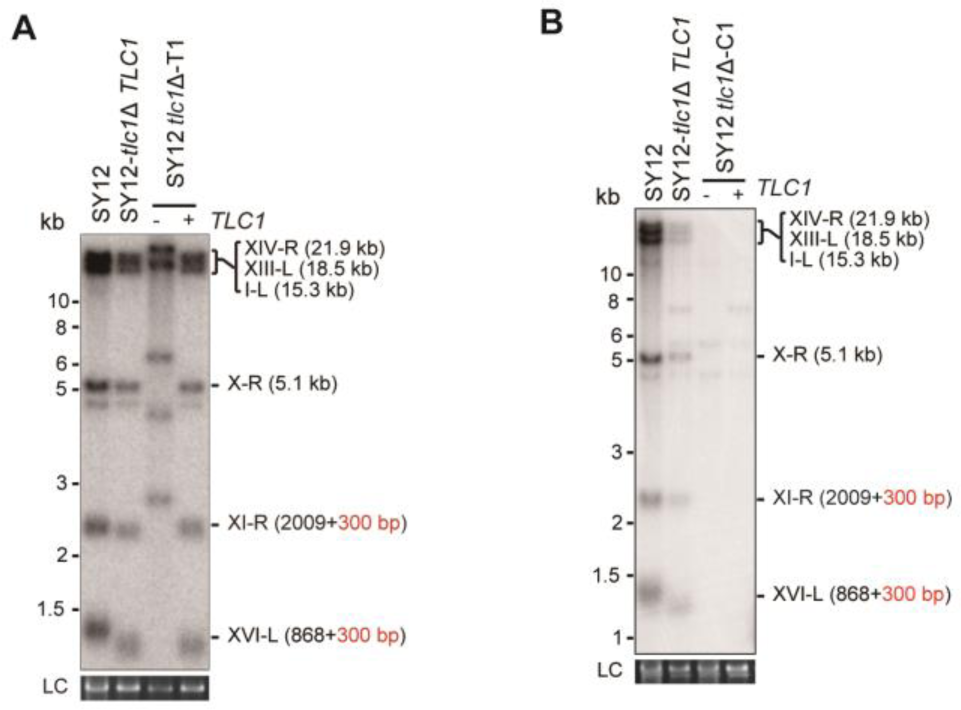
Southern blotting results of reintroduce *TLC1* into SY12 *tlc1*Δ survivors. (**A**) Southern blotting result of SY12 *tlc1*Δ Type II survivor at the 20^th^ re-streaks after *TLC1* reintroduction. LC: loading control. (**B**) Southern blotting result of SY12 *tlc1*Δ circular survivor at the 20^th^ re-streaks after *TLC1* reintroduction. LC: loading control.

**Figure 2—figure supplement 4.**
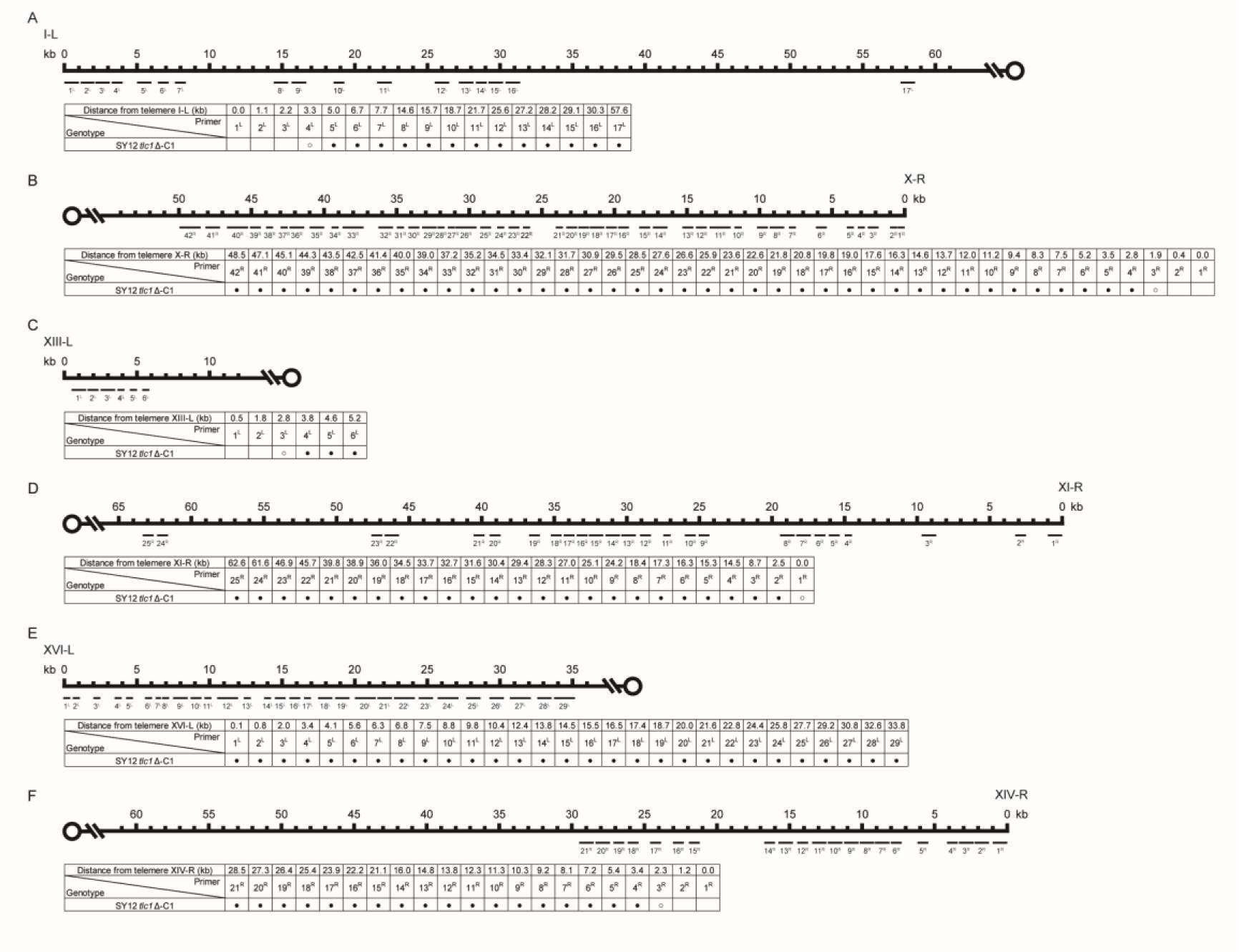
Borders of erosion of the SY12 *tlc1*Δ-C1 survivor are defined by PCR mapping. (**A**) Upper panel, a schematic diagram of the subtelomeric region of 0-60 kb proximal to I-L telomere is shown. Primer pairs (No. 1^L^ to 17^L^) are aligned and indicated at their corresponding subtelomeric loci. Lower panel, the genotype is listed on the left, and primer pairs are listed on top; solid circles mean positive PCR products and open circles mean no PCR products with corresponding primer pairs. (**B**) Upper panel, a schematic diagram of the subtelomeric region of 0-50 kb proximal to X-R telomeric TG1-3 sequence is shown. Primer pairs (No. 1^R^ to 42^R^) are aligned and indicated at their corresponding subtelomeric loci. Lower panel, the genotype is listed on the left, and primer pairs are listed on top. (**C**) Upper panel, a schematic diagram of the subtelomeric region of 0-10 kb proximal to XIII-L telomeric TG1-3 sequence is shown. Primer pairs (No. 1^L^ to 6^L^) are aligned and indicated at their corresponding subtelomeric loci. Lower panel, the genotype is listed on the left, and primer pairs are listed on top. (**D**) Upper panel, a schematic diagram of the subtelomeric region of 0-65 kb proximal to XI-R telomeric TG1-3 sequence is shown. Primer pairs (No. 1^R^ to 25^R^) are aligned and indicated at their corresponding subtelomeric loci. Lower panel, the genotype is listed on the left, and primer pairs are listed on top. (**E**) Upper panel, a schematic diagram of the subtelomeric region of 0-35 kb proximal to XVI-L telomeric TG1-3 sequence is shown. Primer pairs (No. 1^L^ to 29^L^) are aligned and indicated at their corresponding subtelomeric loci. Lower panel, the genotype is listed on the left, and primer pairs are listed on top. (**F**) Upper panel, a schematic diagram of the subtelomeric region of 0-60 kb proximal to XIV-R telomeric TG1-3 sequence is shown. Primer pairs (No. 1^R^ to 21^R^) are aligned and indicated at their corresponding subtelomeric loci. Lower panel, the genotype is listed on the left, and primer pairs are listed on top.

**Figure 2—figure supplement 5.**
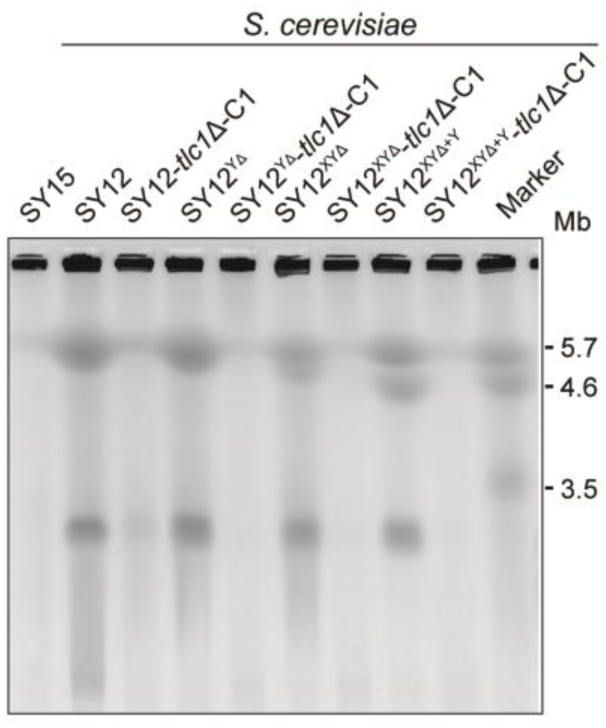
PFGE result of circular survivors. Chromosomal DNA analysis of “circular survivors” SY12 *tlc1*Δ-C1, SY12^YΔ^ *tlc1*Δ-C1, SY12^XYΔ^ *tlc1*Δ-C1, SY12^XYΔ+Y^ *tlc1*Δ-C1 by PFGE. The *S. cerevisiae* strain SY15 (with a single circular chromosome) was used as control. As marker, the size of three chromosomes in wild type *S. pombe* strain is indicated on right.

**Figure 2—figure supplement 6.**
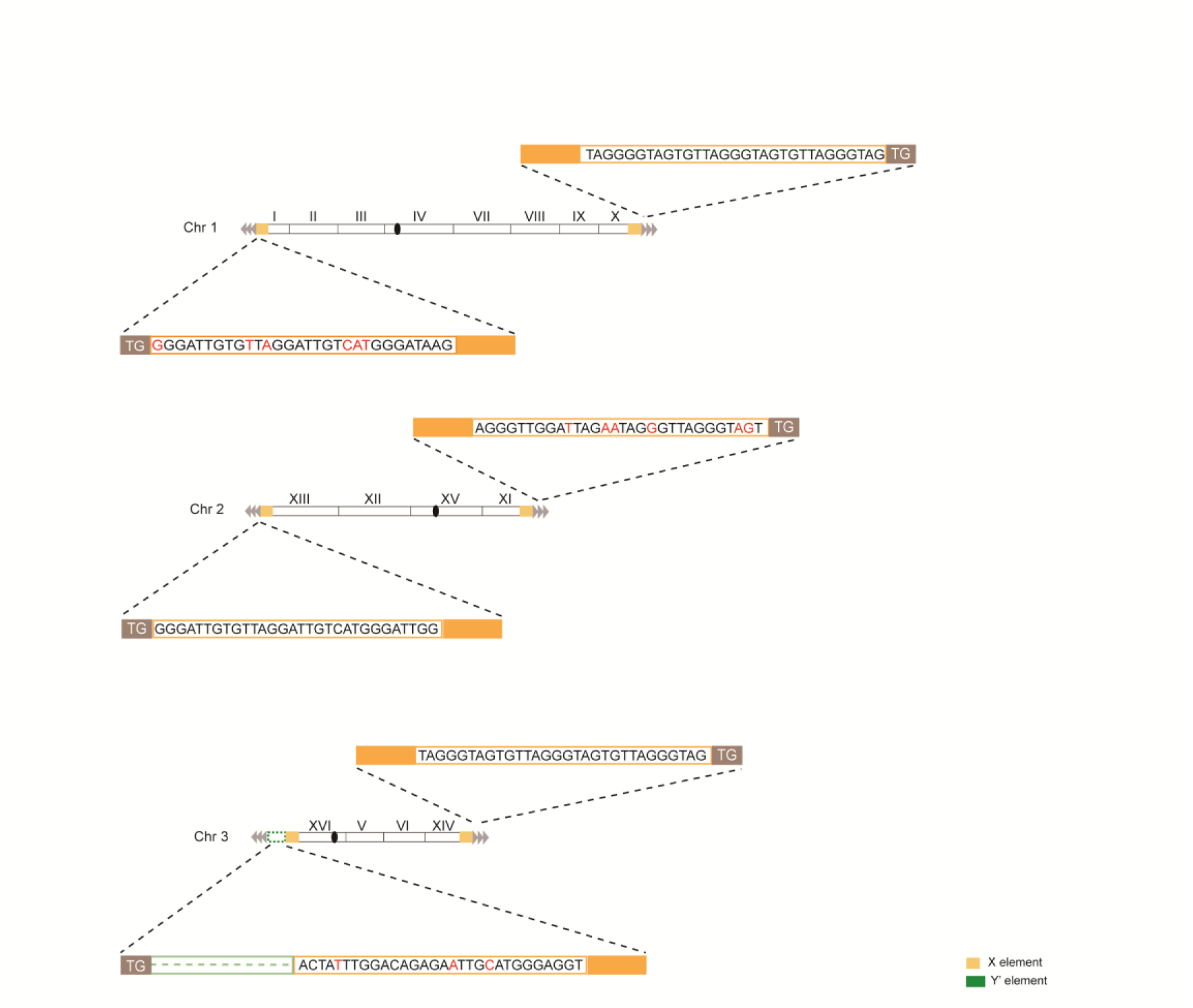
Telomere structure determination of Type X survivor. Schematic representation of chromosome (and telomere) structures (not drawn to scale) in the Type X survivor. The Roman numerals, native chromosomes; the Arabic numerals on the left, chromosome numbers of SY12; yellow box, X-element; green box, Y’-element; tandem grey triangles, telomeres; black circles, centromere. The sequence in yellow box belongs to X-element. Bases in rad are mis-paired. The dotted line in green represents the sequence of Y’-element is lost.

**Figure 2—figure supplement 7.**
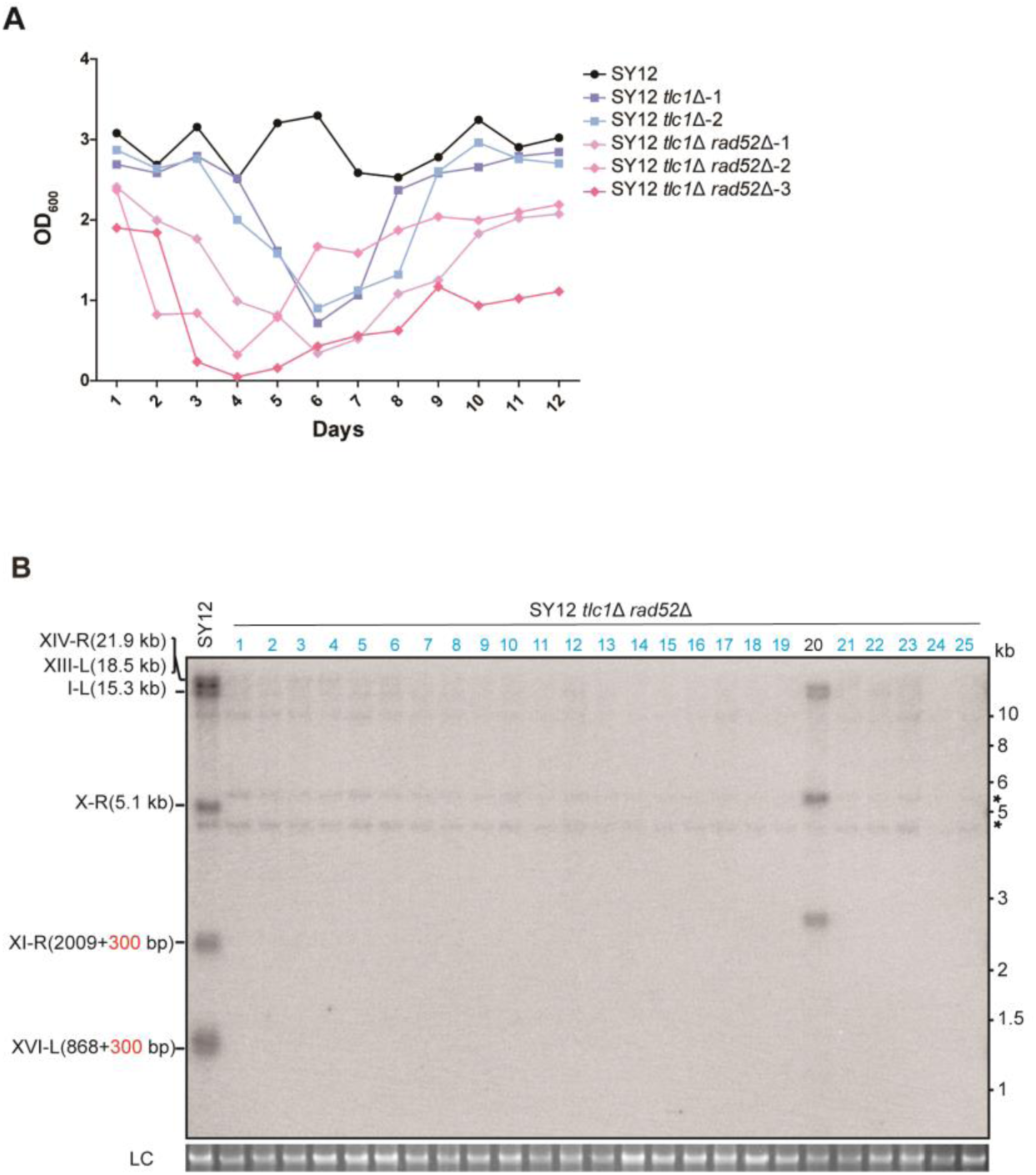
Survivor formation in SY12 *tlc1*Δ *rad52*Δ strain. (**A**) Cell viability assay in liquid culture. The growth of SY12 (labeled in black), SY12 *tlc1*Δ (two clones labeled in purple and blue respectively) and SY12 *tlc1*Δ *rad52*Δ (three clones labeled in violet, pink and carmine respectively) were monitored every 24 hr for 12 days. (**B**) Telomere Southern blotting assay of SY12 *tlc1*Δ *rad52*Δ survivors obtained on solid medium. Genomic DNA from twenty-five independent clones (labeled on top) was digested with XhoI and hybridized to a telomere-specific TG1–3 probe. Circular survivors: in blue; uncharacterized survivors: in black. Theoretical telomere restriction fragments of the SY12 strain are indicated on the left. The asterisks indicate the non-specific bands. Genomic DNA stained with Gelred was used as a relative loading control (LC).

**Figure 2—figure supplement 8.**
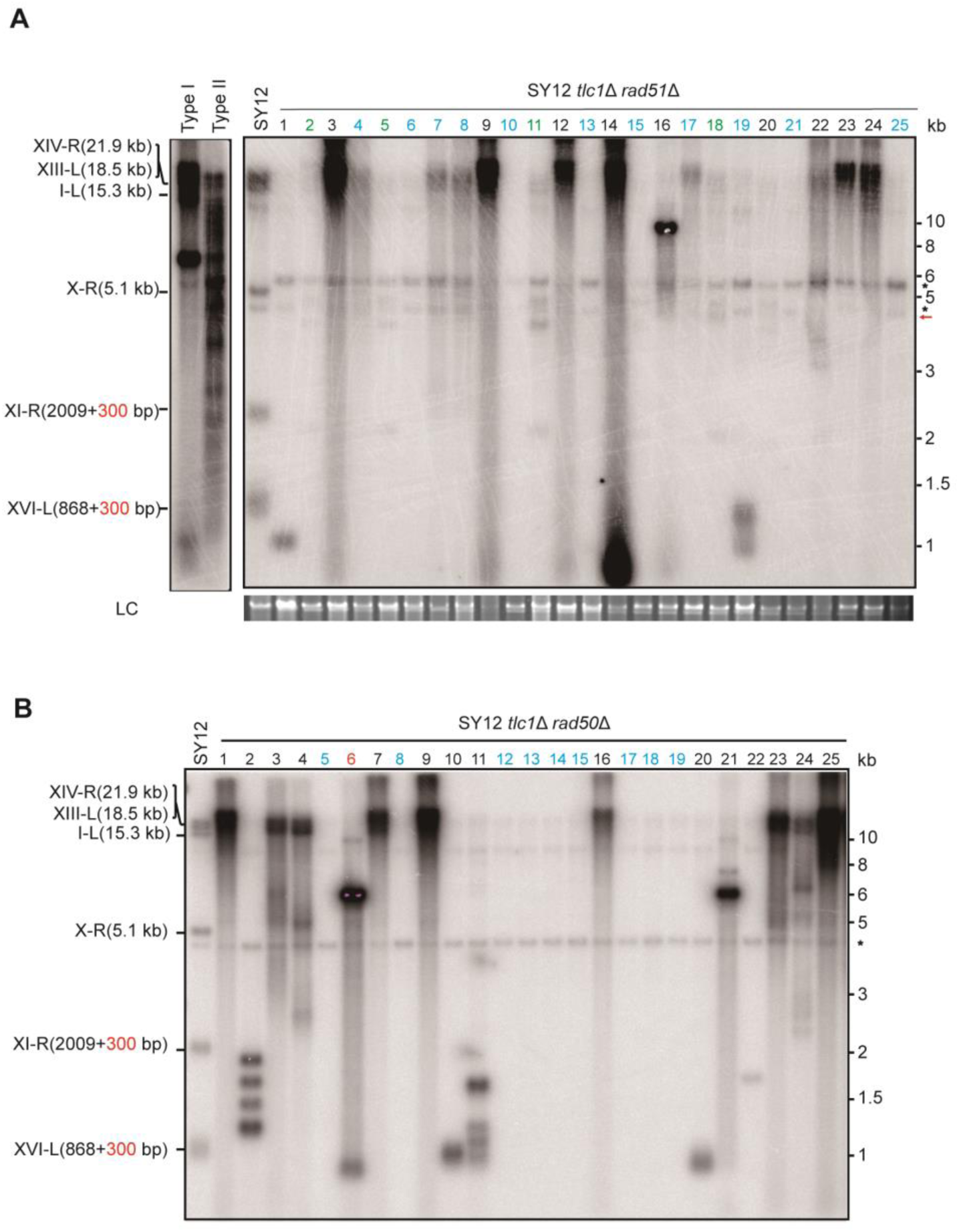
Southern blotting result of SY12 *tlc1*Δ *rad51*Δ and SY12 *tlc1*Δ *rad50*Δ survivors. (**A**) Genomic DNAs of twenty-five independent SY12 *tlc1*Δ *rad51*Δ clones were digested with XhoI and subjected to Southern blotting with a TG1-3 probe. Circular survivors: in blue; Type X survivors: in green; uncharacterized survivors: in black. Theoretical telomere restriction fragments of the SY12 strain are indicated on the left. The red arrows indicate the new band of about 4.3 kb emerged in Type X survivors. The asterisks indicate the non-specific bands. Genomic DNA stained with Gelred was used as a relative loading control (LC). (**B**) Genomic DNAs of twenty-five independent SY12 *tlc1*Δ *rad50*Δ clones were digested with XhoI and subjected to Southern blotting with a TG1-3 probe. Type I survivors: in rad; Circular survivors: in blue; uncharacterized survivors: in black. Theoretical telomere restriction fragments of the SY12 strain are indicated on the left.

**Figure 3—figure supplement 1.**
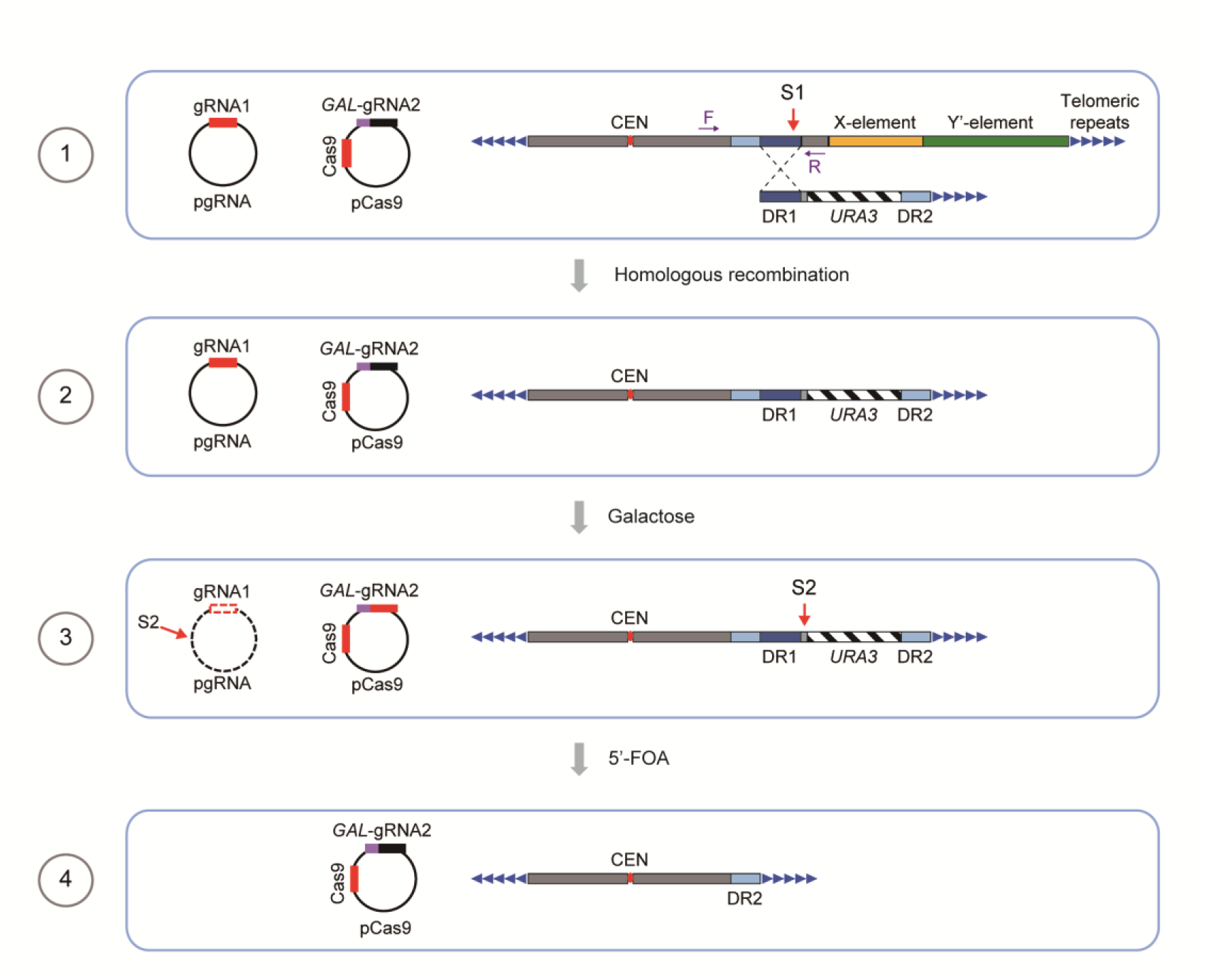
Schematics of CRISPR–Cas9-mediated deletion of X- and Y’-elements on individual chromosomes in SY12 cells. The specific DNA sequences centromere-proximal to the subtelomeric region (site S1) were cleaved by the Cas9 nuclease with the guidance of gRNA1. Homologous recombination between the broken ends and the provided donors led to the deletion of X-and Y’-elements. Galactose induction of gRNA2 on pCas9 caused the cleavage at site S2 and the *URA3* marker was counter-selected on 5’-FOA plates. Deletion of X- and Y’-elements were determined by PCR analysis with a primer located within the deletion region and another upstream of the region (indicated by purple arrows).

**Figure 5—figure supplement 1.**
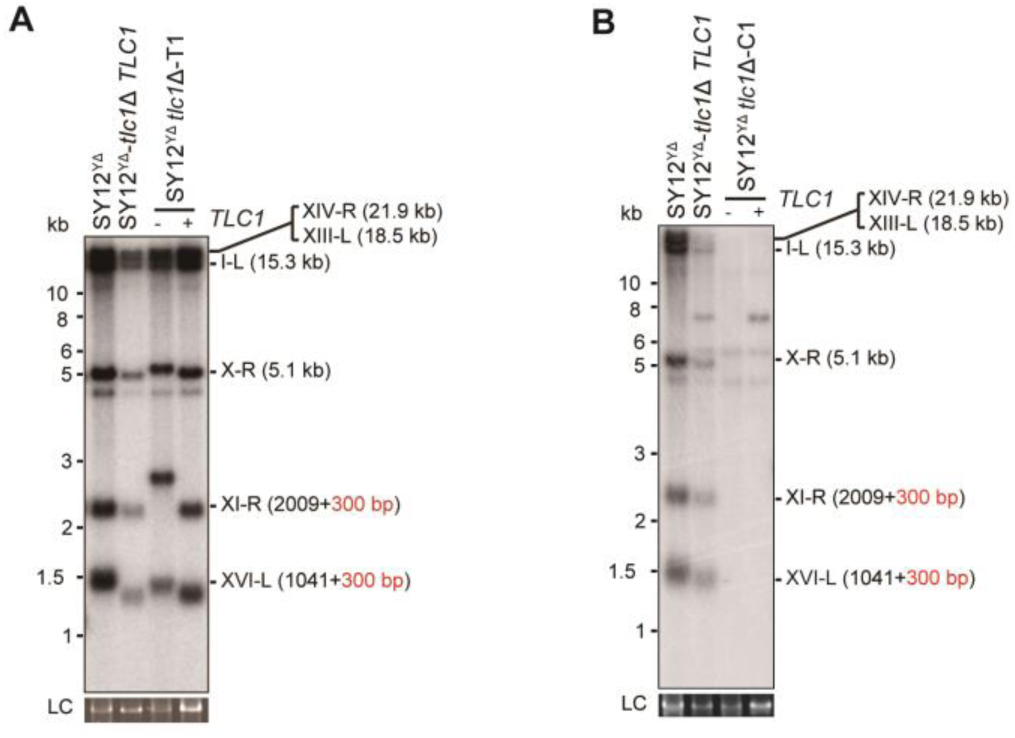
Southern blotting results of reintroducing *TLC1* into SY12 ^YΔ^ *tlc1*Δ survivors. (**A**) Southern blotting result of SY12 ^YΔ^ *tlc1*Δ Type II survivor at the 20^th^ re-streaks after *TLC1* reintroduction. LC: loading control. (**B**) Southern blotting result of SY12 ^YΔ^ *tlc1*Δ circular survivor at the 20^th^ re-streaks after *TLC1* reintroduction. LC: loading control.

**Figure 5—figure supplement 2.**
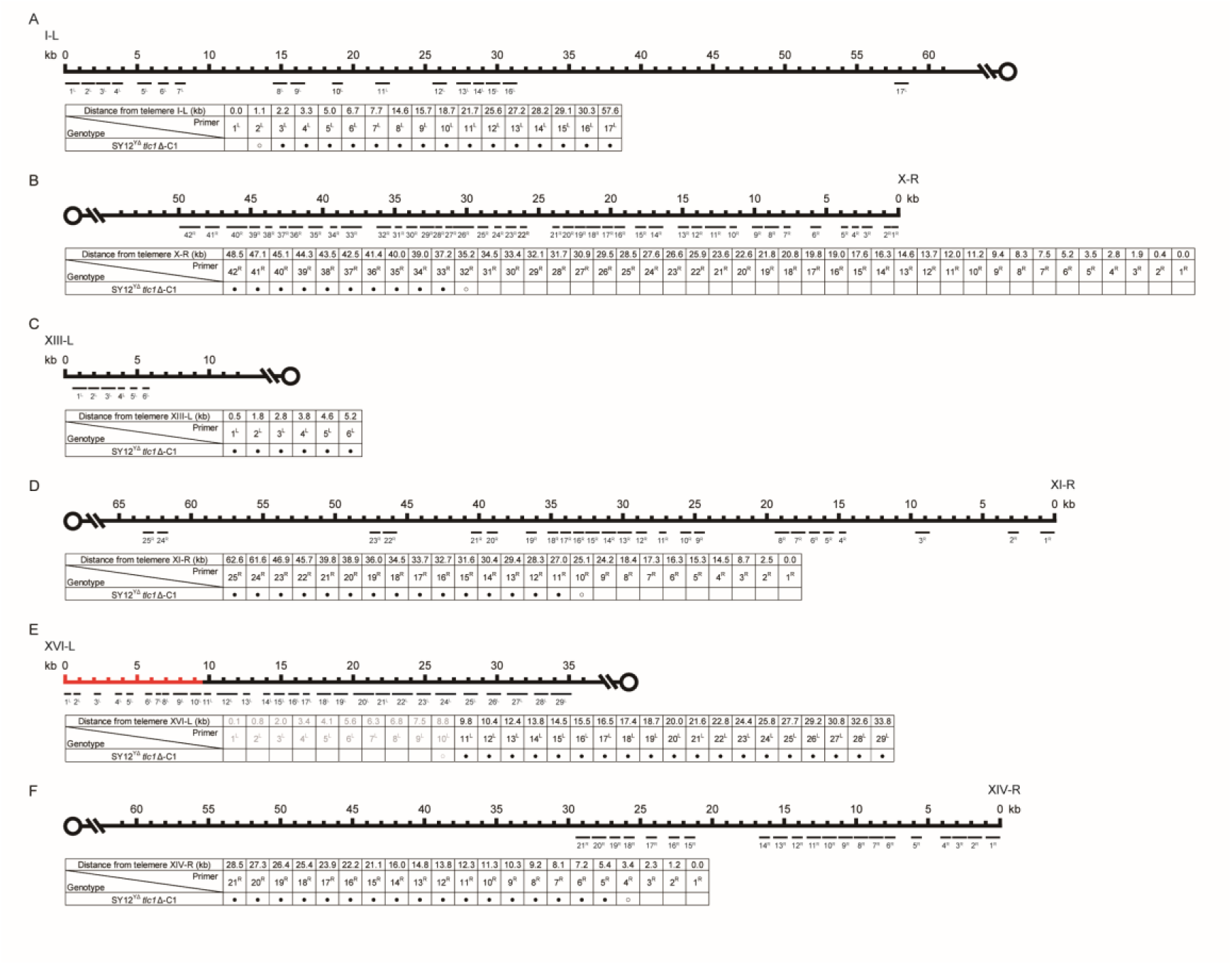
PCR mapping of the borders of erosion in SY12^YΔ^ *tlc1*Δ-C1 cell. Red lines indicate the regions which have been deleted in the SY12^YΔ^ strain.

**Figure 5—figure supplement 3.**
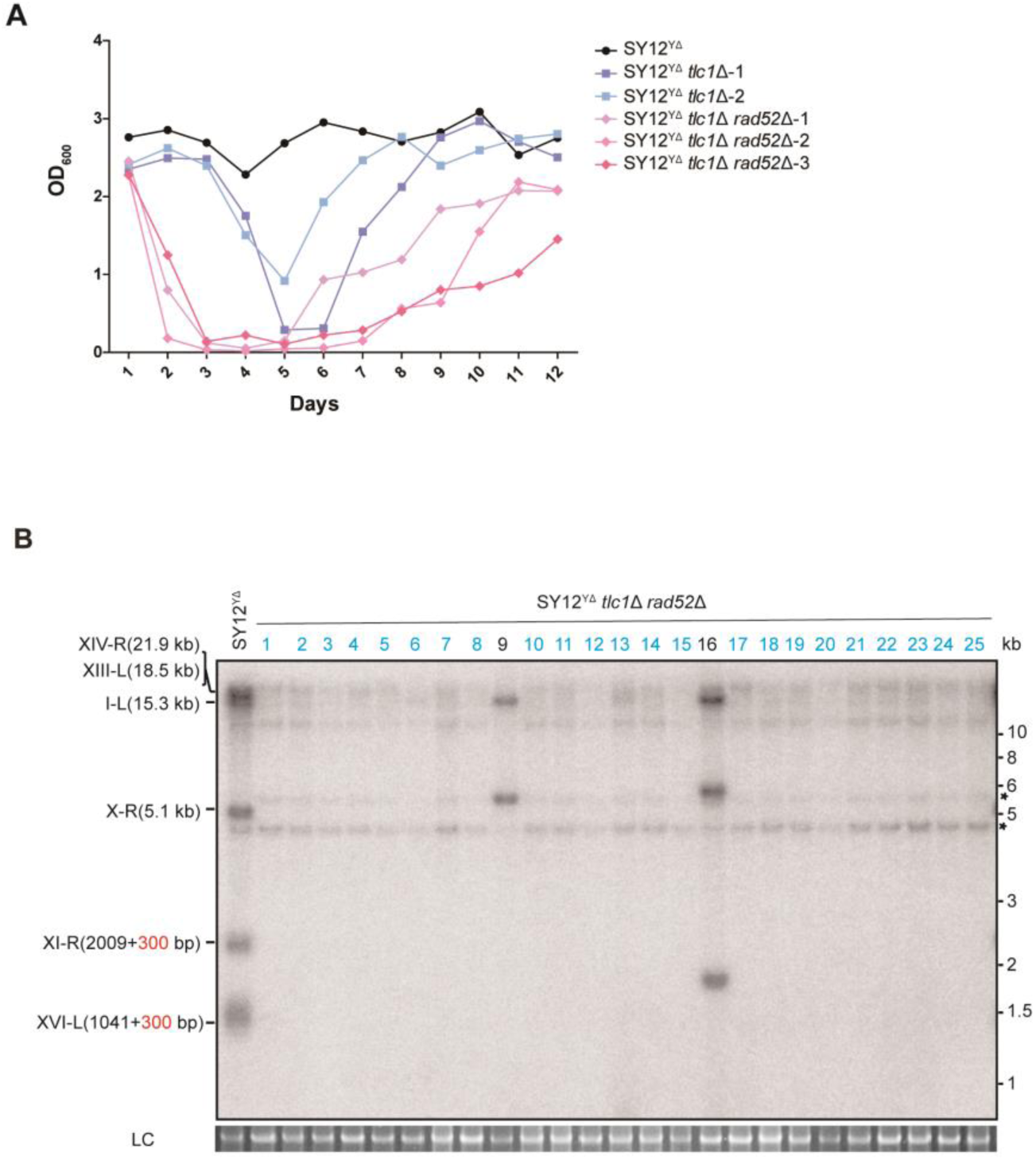
Survivor formation in SY12^YΔ^ *tlc1*Δ *rad52*Δ strain. (**A**) Cell viability assay in liquid culture. The growth of SY12^YΔ^ (labeled in black), SY12^YΔ^ *tlc1*Δ (two clones labeled in purple and blue respectively) and SY12^YΔ^ *tlc1*Δ *rad52*Δ (three clones labeled in violet, pink and carmine respectively) strains were monitored every 24 hr for 12 days. (**B**) Telomere Southern blotting assay of SY12^YΔ^ *tlc1*Δ *rad52*Δ survivors obtained on solid medium. Genomic DNA from twenty-five independent clones (labeled on top) was digested with XhoI and hybridized to a telomere-specific TG1–3 probe. Circular survivors: in blue; uncharacterized survivors: in black. Theoretical telomere restriction fragments of the SY12^YΔ^ strain are indicated on the left. The asterisks indicate the non-specific bands. Genomic DNA stained with Gelred was used as a relative loading control (LC).

**Figure 6—figure supplement 1.**
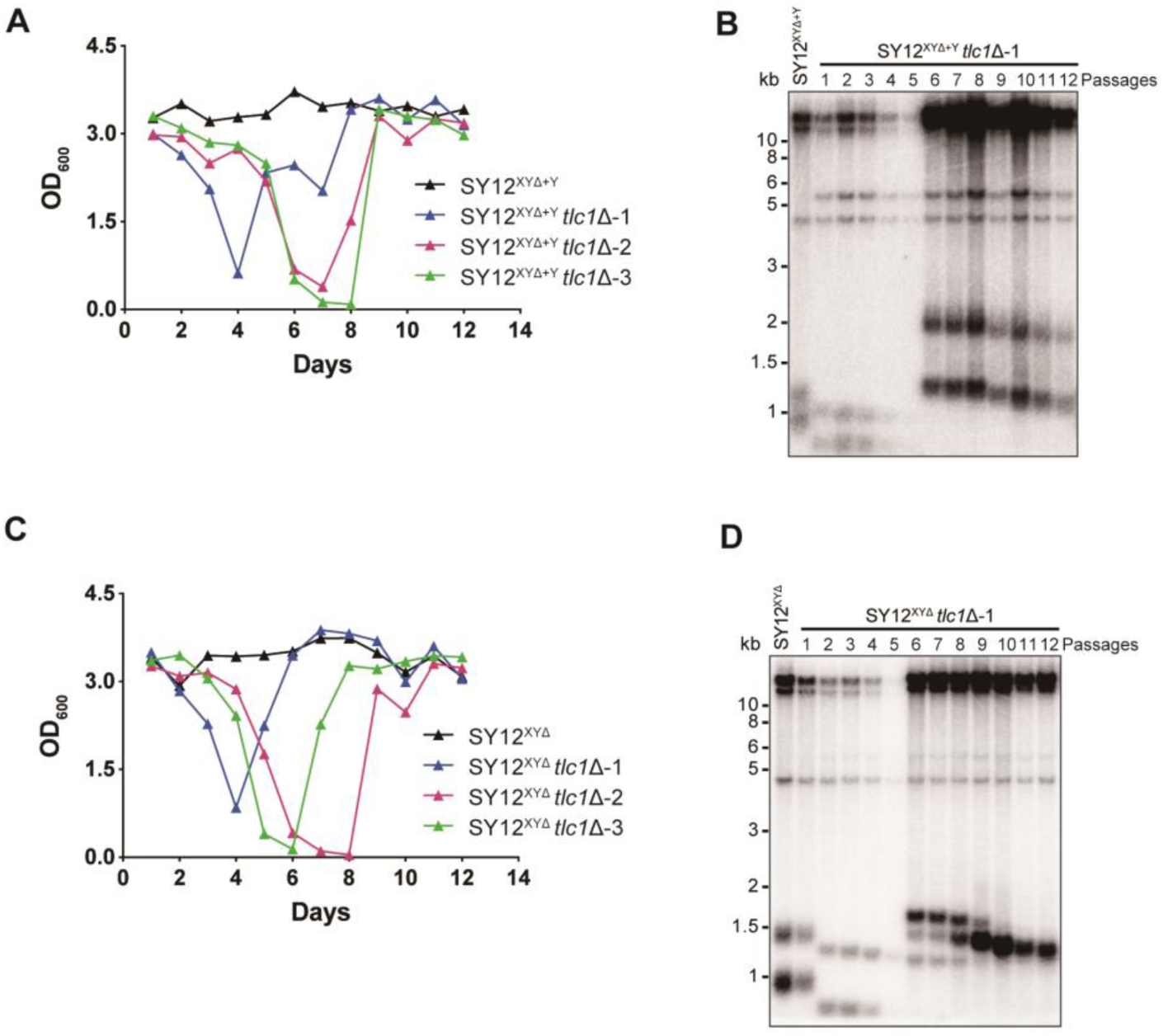
SY12^XYΔ+Y^ *tlc1*Δ and SY12^XYΔ^ *tlc1*Δ strains form Type II survivors in liquid culture. (**A**) Cell viability assay in liquid medium: The growth of SY12^XYΔ+Y^ (labeled in black) and SY12^XYΔ+Y^ *tlc1*Δ (three clones labeled in blue, purple and green respectively) strains were monitored every 24 hr for 12 days. (**B**) Telomeric Southern blotting assay of SY12^XYΔ+Y^ *tlc1*Δ survivors. Genomic DNAs prepared from survivors in (A) were digested with XhoI and subjected to Southern blotting with a TG1-3 probe. (**C**) Cell viability assay in liquid medium: The growth of SY12^XYΔ^ (labeled in black) and SY12^XYΔ^ *tlc1*Δ (three clones labeled in blue, purple and green respectively) strains were monitored every 24 hr for 12 days. (**D**) Telomeric Southern blotting assay of SY12^XYΔ^ *tlc1*Δ survivors. Genomic DNAs prepared from survivors in (C) were digested with XhoI and subjected to Southern blotting with a TG1-3 probe.

**Figure 6—figure supplement 2.**
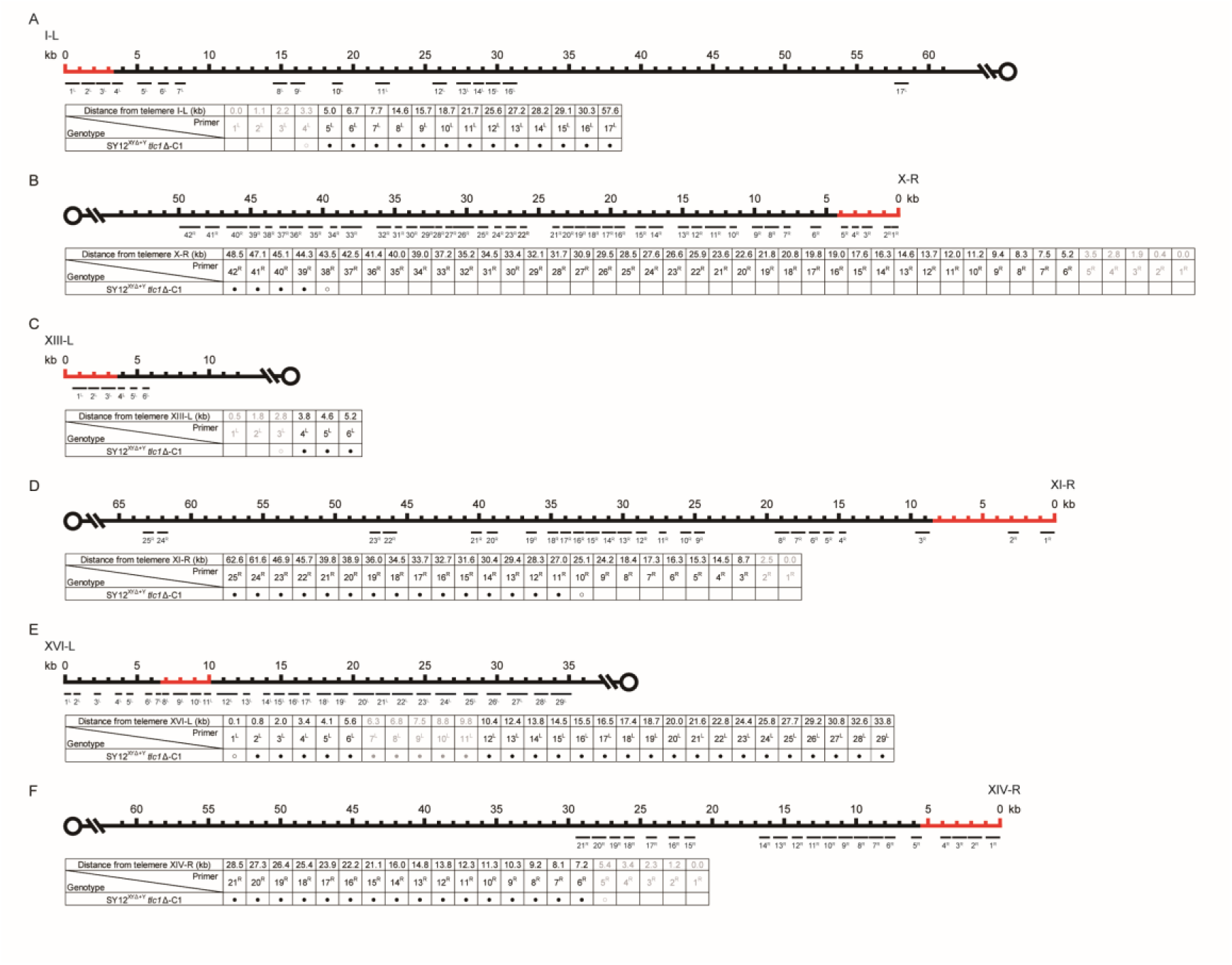
PCR mapping of the borders of erosion in SY12^XYΔ+Υ^ *tlc1*Δ-C1 cell. Red lines indicate the regions which are absent in the SY12^XYΔ+Υ^ strain.

**Figure 6—figure supplement 3.**
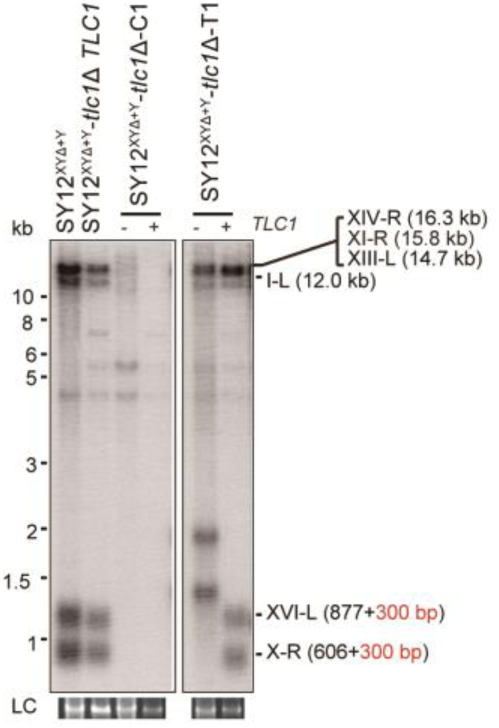
Southern blotting results of an SY12^XYΔ+Y^ *tlc1*Δ circular survivor and an SY12^XYΔ+Y^ *tlc1*Δ Type II survivor at the 20^th^ re-streaks after *TLC1* reintroduction. LC: loading control.

**Figure 6—figure supplement 4.**
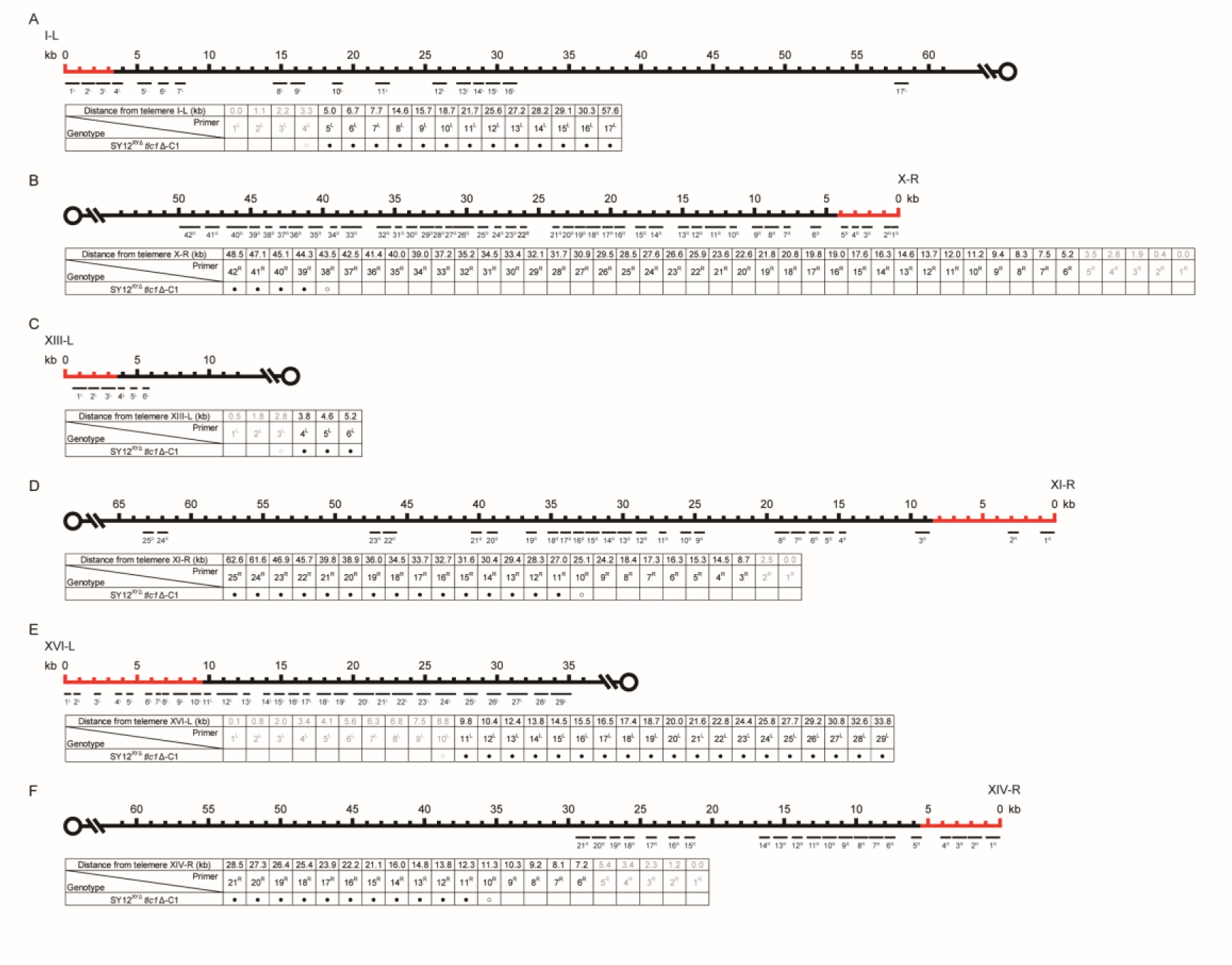
PCR mapping of the borders of erosion in SY12^XYΔ^ *tlc1*Δ-C1 cell. Red lines indicate the regions which have been deleted in the SY12^XYΔ^ strain.

**Figure 6—figure supplement 5.**
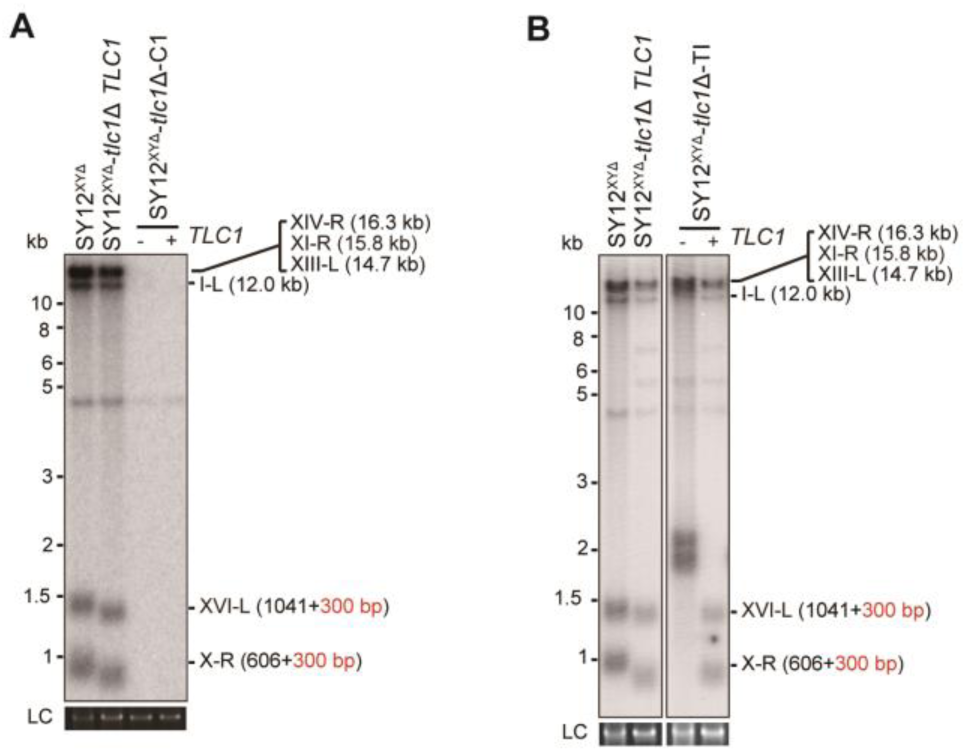
Southern blotting results of reintroducing *TLC1* into SY12^XYΔ^ *tlc1*Δ survivors. (**A**) Southern blotting result of SY12^XYΔ^ *tlc1*Δ circular survivor at the 20^th^ re-streaks after *TLC1* reintroduction. LC: loading control. (**B**) Southern blotting result of SY12^XYΔ^ *tlc1*Δ Type II survivor at the 20^th^ re-streaks after *TLC1* reintroduction. LC: loading control.

**Figure 6—figure supplement 6.**
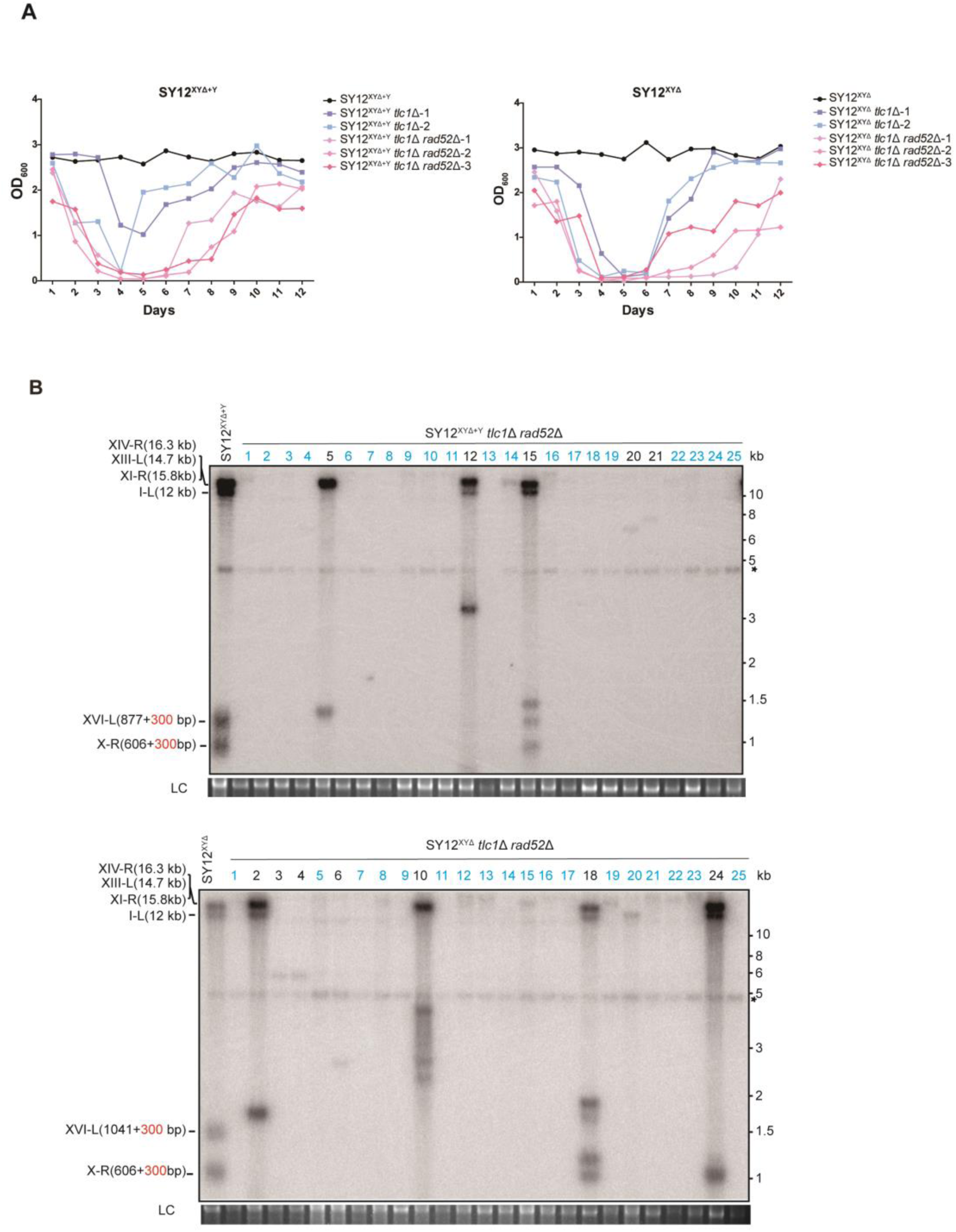
Survivor formation in SY12^XYΔ+Y^ *tlc1*Δ *rad52*Δ and SY12^XYΔ^ *tlc1*Δ *rad52*Δ strains. (**A**) Cell viability assay in liquid culture. The growth of SY12^XYΔ+Y^ (labeled in black), SY12^XYΔ+Y^ *tlc1*Δ (two clones labeled in purple and blue respectively) and SY12^XYΔ+Y^ *tlc1*Δ *rad52*Δ (three clones labeled in violet, pink and carmine respectively) strains shown in the left panel; the survivor formation in SY12^XYΔ^ (labeled in black), SY12^XYΔ^ *tlc1*Δ (two clones labeled in purple and blue respectively) and SY12^XYΔ^ *tlc1*Δ *rad52*Δ (three clones labeled in violet, pink and carmine respectively) strains shown in the right panel. They were monitored every 24 hr for 12 days. (**B**) Telomere Southern blotting assay of SY12^XYΔ+Y^ *tlc1*Δ *rad52*Δ (upper) and SY12^XYΔ^ *tlc1*Δ *rad52*Δ (lower) survivors obtained on solid medium. Genomic DNA from twenty-five independent clones (labeled on top) was digested with XhoI and hybridized to a telomere-specific TG1–3 probe. Circular survivors: in blue; uncharacterized survivors: in black. Theoretical telomere restriction fragments of each strain are indicated on the left. The asterisks indicate the non-specific bands. Genomic DNA stained with Gelred was used as a relative loading control (LC).

**Figure 6—figure supplement 7.**
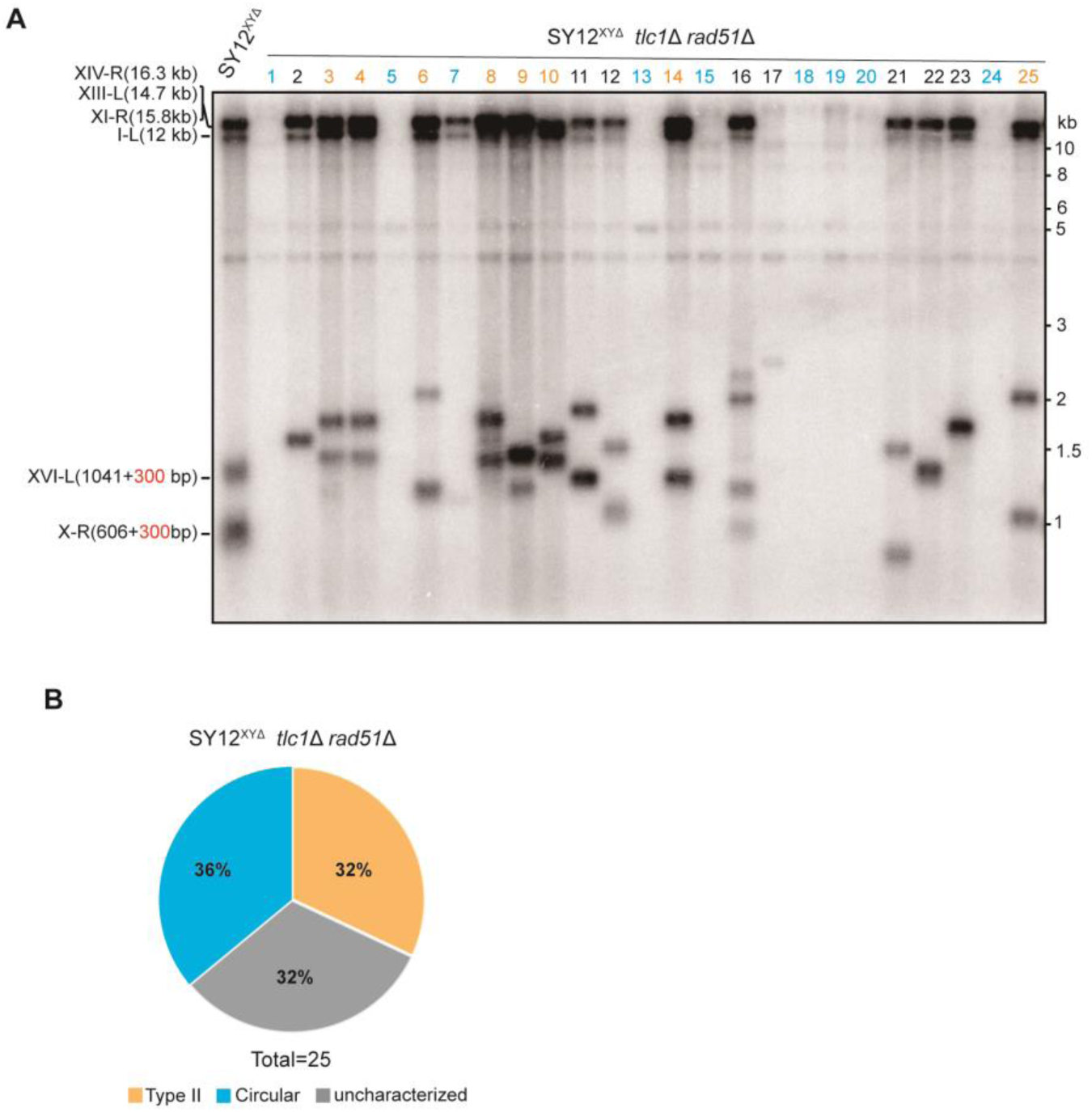
Survivor formation in SY12^XYΔ^ *tlc1*Δ *rad51*Δ strain. (**A**) Southern blotting result of SY12^XYΔ^ *tlc1*Δ *rad51*Δ strain. Twenty-five independent survivors (labeled 1 to 25 on top) were randomly picked, and their genomic DNAs were digested with XhoI and subjected to the Southern blotting assay with a TG1-3 probe. Type II survivors: in orange; circular survivors: in blue; uncharacterized survivors: in black. (**B**) The percentage of survivor types in SY12^XYΔ^ *tlc1*Δ *rad51*Δ strain. n=25; Type II, in orange; uncharacterized survivor, in grey; circular survivor, in blue.

