## Supplementary file 1 for "Elimination of subtelomeric repeat sequences exerts little effect on telomere essential functions in *Saccharomyces cerevisiae*"

| Supplementary file 1. Primers used in this study | | |
| --- | --- | --- |
| Primer name | Sequence | Experiment |
| Telomere-I-L |  |  |
| 1^L^-F | TCCTAACACTACCCTAACAC | PCR-mapping assay |
| 1^L^-R | ACTAGATATAGTCCCTTCGT | PCR-mapping assay |
| 2^L^-F | CGATACTGTGATAGGTACGT | PCR-mapping assay |
| 2^L^-R | AATTAACTTCAATCGCCGCT | PCR-mapping assay |
| 3^L^-F | AAGGCATATAGTTGAAGCAG | PCR-mapping assay |
| 3^L^-R | GCTCTTCAATGAAGGTTTAC | PCR-mapping assay |
| 4^L^-F | TAGAATCTCCCATGTCAACG | PCR-mapping assay |
| 4^L^-R | CTTGTCATGTAGTGGCTTGT | PCR-mapping assay |
| 5^L^-F | CTTGTTCTACTGACAGGATG | PCR-mapping assay |
| 5^L^-R | GAATATCACATCTGCACGTC | PCR-mapping assay |
| 6^L^-F | AAGGAGCTAACAATAGTGGG | PCR-mapping assay |
| 6^L^-R | AAATTACACTCTCGTTGGTT | PCR-mapping assay |
| 7^L^-F | CGGAGTAATCATGGATAACT | PCR-mapping assay |
| 7^L^-R | GATAGGCCATGGTGGAGATT | PCR-mapping assay |
| 8^L^-F | GCTTCATCACAAGAGGATTA | PCR-mapping assay |
| 8^L^-R | GTGAATCCACGAATTACAGC | PCR-mapping assay |
| 9^L^-F | CAGCAACATGGATTGAAAGG | PCR-mapping assay |
| 9^L^-R | CATTAGGACACATTCCTCAT | PCR-mapping assay |
| 10^L^-F | GCCTTACGATATGATGCACA | PCR-mapping assay |
| 10^L^-R | AGTTATATTGCCAACCACAT | PCR-mapping assay |
| 11^L^-F | GTCGAGACGCTTAAAGTTAA | PCR-mapping assay |
| 11^L^-R | ATCAGGGCCAGATTCTCTTC | PCR-mapping assay |
| 12^L^-F | CTGATTAGACCTTCACTAGT | PCR-mapping assay |
| 12^L^-R | CCAGCCAAACTGTACCATTC | PCR-mapping assay |
| 13^L^-F | GTCATCACTAACGGTAGTGC | PCR-mapping assay |
| 13^L^-R | GTTCGCAGTCAGATCAATTA | PCR-mapping assay |
| 14^L^-F | ATGCCATCTGAATTGTCAGG | PCR-mapping assay |
| 14^L^-R | CGAATTACCTCAAAGCCATT | PCR-mapping assay |
| 15^L^-F | GGCATTTCAGCAAGGTAACT | PCR-mapping assay |
| 15^L^-R | AAGATATCGCACTTATTGAA | PCR-mapping assay |
| 16^L^-F | GACGTGCGGTTATATGATCT | PCR-mapping assay |
| 16^L^-R | GCTCAAAGTCAGAAGTCATT | PCR-mapping assay |
| 17^L^-F | TACAACGATTCTCACAAGCA | PCR-mapping assay |
| 17^L^-R | TATACGCATGTGTAGGGTCT | PCR-mapping assay |
| Telomere-X-R |  |  |
| 1^R^-F | TAGCATCCGTGTGCATATGC | PCR-mapping assay |
| 1^R^-R | ACACCACACTACCCTAACAC | PCR-mapping assay |
| 2^R^-F | GGGTGAAGTAAAAGCGTCTG | PCR-mapping assay |
| 2^R^-R | ATTCCACTCCATCACCCATC | PCR-mapping assay |
| 3^R^-F | GGGACAATTGCGCTTCTTTA | PCR-mapping assay |
| 3^R^-R | GACTTTCTCGTAAGCGTTCC | PCR-mapping assay |
| 4^R^-F | CGTTATGCCCGAAGTGTATC | PCR-mapping assay |
| 4^R^-R | ATGAGAACGCCCTTCTGGAC | PCR-mapping assay |
| 5^R^-F | GGAAGTCACTGACGAGGGTT | PCR-mapping assay |
| 5^R^-R | CGGGAGCTACCGTTGAAAAG | PCR-mapping assay |
| 6^R^-F | CGCTCTTGTATCCGGACTGAAC | PCR-mapping assay |
| 6^R^-R | CAGCCAGAACCTGTCCGTAAAC | PCR-mapping assay |
| 7^R^-F | GATGTACGAAGTGAGTGCCCAG | PCR-mapping assay |
| 7^R^-R | GGCTGTGGTTGACCTACCAGAA | PCR-mapping assay |
| 8^R^-F | GTTGCTATTGAACCTGGTGT | PCR-mapping assay |
| 8^R^-R | TATACGGGAGAGTTGCTCTC | PCR-mapping assay |
| 9^R^-F | GGTAGCAGGCATGAAGGAATCG | PCR-mapping assay |
| 9^R^-R | CCACAACCTGTCCGCTTGATTC | PCR-mapping assay |
| 10^R^-F | CCTCGAAAGGTGCAGGTAATGC | PCR-mapping assay |
| 10^R^-R | CCCATCTTCATCACCACTCCGT | PCR-mapping assay |
| 11^R^-F | CGCAGAGGCACAATTTAGCA | PCR-mapping assay |
| 11^R^-R | GCCATCATTGAAGCCGCTCC | PCR-mapping assay |
| 12^R^-F | AGACTGTAACGCCTTGTTGC | PCR-mapping assay |
| 12^R^-R | GCAGGCTAGCTGCATATTCA | PCR-mapping assay |
| 13^R^-F | GTGACTACACCTATGCCTAAG | PCR-mapping assay |
| 13^R^-R | ATGTGTGTTCATGGCTCTTCT | PCR-mapping assay |
| 14^R^-F | GCCAAGATTAGCTTCGAAGA | PCR-mapping assay |
| 14^R^-R | CTTAAAGGGCAGTGGTATCA | PCR-mapping assay |
| 15^R^-F | GTGCTCTTCGAAATGTCTGA | PCR-mapping assay |
| 15^R^-R | TCGCTTTAGACCGAACATAA | PCR-mapping assay |
| 16^R^-F | GAAGATCTTGGCGTATGCCA | PCR-mapping assay |
| 16^R^-R | TTGCAAGGCTTCCAATGGTA | PCR-mapping assay |
| 17^R^-F | TAACTGTGAGTTGGCGTACA | PCR-mapping assay |
| 17^R^-R | CCACCTATATTGTATGGCTC | PCR-mapping assay |
| 18^R^-F | GATGTTCTGGAATCCCATCT | PCR-mapping assay |
| 18^R^-R | AATTGTACTCACGATGGGAG | PCR-mapping assay |
| 19^R^-F | GTTTGAAGGCACAACCACAT | PCR-mapping assay |
| 19^R^-R | GAAGAACCACCGGTAATTGA | PCR-mapping assay |
| 20^R^-F | CAATTGGCAACTGAAAGCTA | PCR-mapping assay |
| 20^R^-R | GTAACAGTAGTACCACTGCT | PCR-mapping assay |
| 21^R^-F | GTCGACTTGTCCTGCCTCATAC | PCR-mapping assay |
| 21^R^-R | CATGGCCGTGCTAGCAGTAACA | PCR-mapping assay |
| 22^R^-F | CTCTGGAGTGTCCTTTCCCAGT | PCR-mapping assay |
| 22^R^-R | GTACCGTGCTTAGAACTGGCTC | PCR-mapping assay |
| 23^R^-F | CATCCACTTCCCCATAGTGC | PCR-mapping assay |
| 23^R^-R | CCACCATGGATATTGTGCTG | PCR-mapping assay |
| 24^R^-F | GCCCTACATGCACAACAAAT | PCR-mapping assay |
| 24^R^-R | GGCCAACGCCGTATACTAAC | PCR-mapping assay |
| 25^R^-F | CAGTAGACACTGGGTCACTTGG | PCR-mapping assay |
| 25^R^-R | AATGGAAGACATATCGGCCTACGG | PCR-mapping assay |
| 26^R^-F | GCGGTAGAAATGGTAGAAGT | PCR-mapping assay |
| 26^R^-R | GTGAATGTCTATTGAACGACG | PCR-mapping assay |
| 27^R^-F | CACTGAAAGTAGACCCGAAG | PCR-mapping assay |
| 27^R^-R | CTAGTGCCTCTGCATCCTCT | PCR-mapping assay |
| 28^R^-F | AGTAGACGACTCTGGAGAGGAAC | PCR-mapping assay |
| 28^R^-R | CAGTAAAGCAACCACTTCCGCAG | PCR-mapping assay |
| 29^R^-F | AGCCGAGGAGGCTTTTGGAA | PCR-mapping assay |
| 29^R^-R | CTGAGCGGACTTCTTCCTTA | PCR-mapping assay |
| 30^R^-F | CATTCGTCATCGCCGCATCA | PCR-mapping assay |
| 30^R^-R | GGTACCGCTATCGTTGCTGT | PCR-mapping assay |
| 31^R^-F | GCTACGACTGTTGAAATCGT | PCR-mapping assay |
| 31^R^-R | ACCCGTTCCGACAACGACAA | PCR-mapping assay |
| 32^R^-F | AGGTGGAAGAAGCCTCTGTGGT | PCR-mapping assay |
| 32^R^-R | AAGGGCAAACAGGCCTGAGGTA | PCR-mapping assay |
| 33^R^-F | CTGGCCTATCGGTATCAAGGAC | PCR-mapping assay |
| 33^R^-R | CACATCCTGTGAACGGTTACGC | PCR-mapping assay |
| 34^R^-F | GTCTTTAGACCCTTCCGCGGTG | PCR-mapping assay |
| 34^R^-R | CAATCGCAGCAGTACCAGAACC | PCR-mapping assay |
| 35^R^-F | GACAGGCTTCAGGGCAATCT | PCR-mapping assay |
| 35^R^-R | GGGTGCCAAGGTCATATCGT | PCR-mapping assay |
| 36^R^-F | GCCGCTGCTACTTTCAACTG | PCR-mapping assay |
| 36^R^-R | TCGCTCTAGGATGACTTTGG | PCR-mapping assay |
| 37^R^-F | CGGTTCTAACCTTACCGTCCAC | PCR-mapping assay |
| 37^R^-R | GGCAAGTCCTGTTCTGTGTGGT | PCR-mapping assay |
| 38^R^-F | AAGATCTCGGCGTCGGTTCTGA | PCR-mapping assay |
| 38^R^-R | CTGGTGGTCGTAACTTGGGTCG | PCR-mapping assay |
| 39^R^-F | GGGCACAGACATCTGTACTTCTG | PCR-mapping assay |
| 39^R^-R | CCCTAGTGACCAGCTTGGATGT | PCR-mapping assay |
| 40^R^-F | CGTCACCGTTGGTCCAGAACGA | PCR-mapping assay |
| 40^R^-R | CCACCTTCCACTAAGGACGTAGA | PCR-mapping assay |
| 41^R^-F | TGAGCGTGCAGTAGCAGGTGTT | PCR-mapping assay |
| 41^R^-R | TCGCGTGCTTATTCTCAGGAGC | PCR-mapping assay |
| 42^R^-F | GACCTGCCTTGCTACCGTCTAT | PCR-mapping assay |
| 42^R^-R | AGGGTCTACGCGGTCCATAATC | PCR-mapping assay |
| Telomere-XIII-L |  |  |
| 1^L^-F | ACTCCCACGATTATCCACAT | PCR-mapping assay |
| 1^L^-R | TCGGTAGTGAGATGGCAGTT | PCR-mapping assay |
| 2^L^-F | TCTGGTGGAAACTTTCCAAC | PCR-mapping assay |
| 2^L^-R | GGGATAATTGCGCTTCTTTA | PCR-mapping assay |
| 3^L^-F | GCAGTGAGGTGCTCTTAGTG | PCR-mapping assay |
| 3^L^-R | CCAAAGTGTGAACGAAGTTG | PCR-mapping assay |
| 4^L^-F | ACAACATTACCAGTCACTTC | PCR-mapping assay |
| 4^L^-R | CTAGTCTTAGACCTATGCCT | PCR-mapping assay |
| 5^L^-F | AGGCATAGGTCTAAGACTAG | PCR-mapping assay |
| 5^L^-R | ACACCGACTACATTATCACC | PCR-mapping assay |
| 6^L^-F | GGTGATAATGTAGTCGGTGT | PCR-mapping assay |
| 6^L^-R | GAGCAACACAGTTTATCTTA | PCR-mapping assay |
| Telomere-XI-R |  |  |
| 1^R^-F | GAATGTGGTAACCCAATAGC | PCR-mapping assay |
| 1^R^-R | AACCCTAACACTACCCTACT | PCR-mapping assay |
| 2^R^-F | ATAGACAAGGCGACACACGA | PCR-mapping assay |
| 2^R^-R | TATCGCGGAGGTAGCTATCA | PCR-mapping assay |
| 3^R^-F | CATATCACCTCCAGCTAATT | PCR-mapping assay |
| 3^R^-R | ACAGACAGGTTCAAAGGAGT | PCR-mapping assay |
| 4^R^-F | AAGTAGGATAGGCATTCTTT | PCR-mapping assay |
| 4^R^-R | TGGTAATCCCAGAACCTATT | PCR-mapping assay |
| 5^R^-F | ACAATAGTTACTACTGCCCG | PCR-mapping assay |
| 5^R^-R | GACGACGTAGAGCTCTCTAC | PCR-mapping assay |
| 6^R^-F | AGTCTGGTGTCTGTTCCGAA | PCR-mapping assay |
| 6^R^-R | AAGCTTCAGCTGTTGGTTGA | PCR-mapping assay |
| 7^R^-F | ACCAACGCTCCAACTTCATC | PCR-mapping assay |
| 7^R^-R | ACCGTTGTTAATTCCGTAGC | PCR-mapping assay |
| 8^R^-F | GACGATAATGCTACTCAAGC | PCR-mapping assay |
| 8^R^-R | TGAAGTAGCTGTGCCAGTAG | PCR-mapping assay |
| 9^R^-F | CTAGCGAGGTCCTAGATCAA | PCR-mapping assay |
| 9^R^-R | AATGTCGCATCCGTCTAATG | PCR-mapping assay |
| 10^R^-F | GCAGTTATTGATGGATGGCT | PCR-mapping assay |
| 10^R^-R | TGCGTATGCACATTCTCTAT | PCR-mapping assay |
| 11^R^-F | CCATCATCCATGAAGAGTTC | PCR-mapping assay |
| 11^R^-R | AACTACGGCAATAATGCCAG | PCR-mapping assay |
| 12^R^-F | TGGACAACATGGATAGCGAT | PCR-mapping assay |
| 12^R^-R | GGATTCAGTGGCAGGACTGA | PCR-mapping assay |
| 13^R^-F | ATGAGCAGCTCTTGAAAGCT | PCR-mapping assay |
| 13^R^-R | TATTGCTGCTGACAGGATGG | PCR-mapping assay |
| 14^R^-F | AGCAGCATGGTCTAACAATC | PCR-mapping assay |
| 14^R^-R | GTCCAGAGATCCTGTCCAAC | PCR-mapping assay |
| 15^R^-F | TTCGTTCGAGAGCTCTTCAT | PCR-mapping assay |
| 15^R^-R | CAATCAGGAAATGGAAGCTT | PCR-mapping assay |
| 16^R^-F | CCGATGCAATTAACAGAATT | PCR-mapping assay |
| 16^R^-R | CTAGTGTCATATGATGCGAA | PCR-mapping assay |
| 17^R^-F | ATCTATGACGAGAAGTCGCA | PCR-mapping assay |
| 17^R^-R | GCTTGTTGATCCAGTTTCAG | PCR-mapping assay |
| 18^R^-F | GTAGGATCTACTTCCGAAGT | PCR-mapping assay |
| 18^R^-R | CCAAACTTGATAGCGTCGAA | PCR-mapping assay |
| 19^R^-F | GAACTAAGTGTCAACGAGGG | PCR-mapping assay |
| 19^R^-R | GATATTCTCCGGTGGATCGT | PCR-mapping assay |
| 20^R^-F | TTCGATGTTTCTGTCTGCAT | PCR-mapping assay |
| 20^R^-R | AGAGTTTGTTAGTTGTGAGC | PCR-mapping assay |
| 21^R^-F | TCACTGGATCACACATTGTA | PCR-mapping assay |
| 21^R^-R | CCTTAATGGATTCTGAACAG | PCR-mapping assay |
| 22^R^-F | ATATATTGCTGCCTGTCTATT | PCR-mapping assay |
| 22^R^-R | CTTCATCTGCTTCGTCAGTT | PCR-mapping assay |
| 23^R^-F | TTGAACCTTTGGTGGCTTTA | PCR-mapping assay |
| 23^R^-R | ATTGTAAATGGAGCGTATCT | PCR-mapping assay |
| 24^R^-F | TGAAGGACCATTGTCATAGC | PCR-mapping assay |
| 24^R^-R | TTCCTCCTATGATTGGCTAC | PCR-mapping assay |
| 25^R^-F | TTTATGGGAAGAGCGTGCTA | PCR-mapping assay |
| 25^R^-R | AGCATTCTTCTCGATAAGTC | PCR-mapping assay |
| Telomere-XVI-L |  |  |
| 1^L^-F | CTCATGTACGTCTCCTCCAAGC | PCR-mapping assay |
| 1^L^-R | CTGTCGATGCTGATAGGGCTGT | PCR-mapping assay |
| 2^L^-F | CTCCTCAACTGTCGATGATGCC | PCR-mapping assay |
| 2^L^-R | ATCGAGCCATATCATGGGGACC | PCR-mapping assay |
| 3^L^-F | TCGCATTGGTACTGGCATTAGC | PCR-mapping assay |
| 3^L^-R | TGTCACTGACGGTAGCATGCGA | PCR-mapping assay |
| 4^L^-F | CAGAGGGTGTGTCTGCCATGTA | PCR-mapping assay |
| 4^L^-R | GGCACTGCGTAGGTAGCAGATT | PCR-mapping assay |
| 5^L^-F | CGTCAAGCGTCTTAAGTCGAGGC | PCR-mapping assay |
| 5^L^-R | CACGAAGTAGCGCAGGAAACCG | PCR-mapping assay |
| 6^L^-F | CCGACCATGACGGAAACGACAA | PCR-mapping assay |
| 6^L^-R | GCGGCTGAGCAACGAACAGAAT | PCR-mapping assay |
| 7^L^-F | GCGCTTCGAAGTAGAGGAGCC | PCR-mapping assay |
| 7^L^-R | GGAGAGGTAGGGTAATGGAGGG | PCR-mapping assay |
| 8^L^-F | ACCATCCACCGCCCATCATAAC | PCR-mapping assay |
| 8^L^-R | CTGAAGTGGCATCCGTTCAAGC | PCR-mapping assay |
| 9^L^-F | GCTCACACACATGTGGGCGCTA | PCR-mapping assay |
| 9^L^-R | TGGCATTCTCACATTCGGCGCA | PCR-mapping assay |
| 10^L^-F | CCGTGCCTGTGATACTTCCT | PCR-mapping assay |
| 10^L^-R | GCGTATCGAAGAGGAACTGG | PCR-mapping assay |
| 11^L^-F | TGACCACATCTTAAACCAACG | PCR-mapping assay |
| 11^L^-R | CCCGATATTGACACTGCCGA | PCR-mapping assay |
| 12^L^-F | CCAGCGGAAGGTCCATATTGCT | PCR-mapping assay |
| 12^L^-R | CAGCCAAATCCACCAGTCTCAG | PCR-mapping assay |
| 13^L^-F | GGTGTGATCGCTGCCATCTGTC | PCR-mapping assay |
| 13^L^-R | GTGCTACCGACCTGCCGTTTTC | PCR-mapping assay |
| 14^L^-F | GTGCGTACGCGAGTTTATCCA | PCR-mapping assay |
| 14^L^-R | CGAGCGTGTAATGCTCTGATG | PCR-mapping assay |
| 15^L^-F | GAGCCACCAGACGCTAAATA | PCR-mapping assay |
| 15^L^-R | CGCTCCACTATCGATGGTTT | PCR-mapping assay |
| 16^L^-F | CTCCGACAACGCTGACAGCA | PCR-mapping assay |
| 16^L^-R | TTATAGAGCAGCACGGGACC | PCR-mapping assay |
| 17^L^-F | TGGTGCCATTGCCGAACCTC | PCR-mapping assay |
| 17^L^-R | AAGCAGCGGCAGTGAACTAC | PCR-mapping assay |
| 18^L^-F | GATTCTGGAGGTTCAAATAA | PCR-mapping assay |
| 18^L^-R | TAATTGCTTATTGATGACCA | PCR-mapping assay |
| 19^L^-F | GTACTACGACTACCAGGAAC | PCR-mapping assay |
| 19^L^-R | CGCAACTAGCATACATTTAT | PCR-mapping assay |
| 20^L^-F | TTCTTGCTGTATTCACGAGC | PCR-mapping assay |
| 20^L^-R | TTAGTAGGTCGAGACCAGAA | PCR-mapping assay |
| 21^L^-F | TTCAGCAGTAACACGCTGGA | PCR-mapping assay |
| 21^L^-R | GCGAATATAGCTTGTAACCA | PCR-mapping assay |
| 22^L^-F | TTGCATAGACATCGCTGTCG | PCR-mapping assay |
| 22^L^-R | ACCAAATGCACCACTAATCC | PCR-mapping assay |
| 23^L^-F | ATCATTCTTCACCTGGTTCT | PCR-mapping assay |
| 23^L^-R | TCAATATTTCTGCGCCAGCA | PCR-mapping assay |
| 24^L^-F | GGATGAGAGTATCCTGTCTA | PCR-mapping assay |
| 24^L^-R | TCATCTGTGGTATTGCAATG | PCR-mapping assay |
| 25^L^-F | GAGATTGATGTCATTGACAT | PCR-mapping assay |
| 25^L^-R | AGCGAATCTGACCATTGTAT | PCR-mapping assay |
| 26^L^-F | CGATTGTGAATTCGGATAGC | PCR-mapping assay |
| 26^L^-R | GGTTCAGAAGCTGCAGTGCC | PCR-mapping assay |
| 27^L^-F | CTTCTCTGCCTTCAGCCTCT | PCR-mapping assay |
| 27^L^-R | GGTATATGTCAACCCATAAG | PCR-mapping assay |
| 28^L^-F | CCTTGGCCATGATGGATCAGT | PCR-mapping assay |
| 28^L^-R | TAGTAAGCCATGGCTCCAAC | PCR-mapping assay |
| 29^L^-F | ATTTACCCTAGAACAACTAG | PCR-mapping assay |
| 29^L^-R | TTAATGACCTGAGGCTGCTT | PCR-mapping assay |
| Telomere-XIV-R |  |  |
| 1^R^-F | CGTTTCGTTGACTCCTACAA | PCR-mapping assay |
| 1^R^-R | CTACCCTAACACTACCCTAA | PCR-mapping assay |
| 2^R^-F | GAGCATCTGTTAACGAATCG | PCR-mapping assay |
| 2^R^-R | GTCTCACTTCATCTTACCAC | PCR-mapping assay |
| 3^R^-F | CAGCATGCTTGGTATCTTAT | PCR-mapping assay |
| 3^R^-R | AGCACCTCACATATTCTCCA | PCR-mapping assay |
| 4^R^-F | GATGAACGCGCTGTCAACAA | PCR-mapping assay |
| 4^R^-R | CAGACATATCAGGAAGGAAG | PCR-mapping assay |
| 5^R^-F | CAATCTGCATCAACGAGACT | PCR-mapping assay |
| 5^R^-R | TCACTTGTACAGGGAACTGG | PCR-mapping assay |
| 6^R^-F | TAGTGTCCTTGATACCACGC | PCR-mapping assay |
| 6^R^-R | CCTTCTTATGTGCCTATCTT | PCR-mapping assay |
| 7^R^-F | GGTCAACTGTGCAACGAATT | PCR-mapping assay |
| 7^R^-R | CGATGATCCAAGAGCTGTTA | PCR-mapping assay |
| 8^R^-F | GTGAACTTCTTTTCCACTATTATTGCTGTTATGG | PCR-mapping assay |
| 8^R^-R | AATTCGTTGCACAGTTGACCCAC | PCR-mapping assay |
| 9^R^-F | CGTGGAAAAAGAGTGGTCAGATGGA | PCR-mapping assay |
| 9^R^-R | TAACAGCAATAATAGTGGAAAAGAAGTTCACTGTAC | PCR-mapping assay |
| 10^R^-F | TGGTTCGTAGGCTATTCAGTGGC | PCR-mapping assay |
| 10^R^-R | CATCTGACCACTCTTTTTCCACGG | PCR-mapping assay |
| 11^R^-F | TAGCGTCAATAATGAAAGTGCAATCAAAGAC | PCR-mapping assay |
| 11^R^-R | CCACTGAATAGCCTACGAACCAC | PCR-mapping assay |
| 12^R^-F | TTGTCAGTACAAAGTAGGTG | PCR-mapping assay |
| 12^R^-R | GTAACAGGCGGACTCTGTAG | PCR-mapping assay |
| 13^R^-F | CTCTAGCAGGTCAATCAATT | PCR-mapping assay |
| 13^R^-R | ATCTGCTGTTGCATCTGGAT | PCR-mapping assay |
| 14^R^-F | ATTGTGGTGGCACTTTGGTT | PCR-mapping assay |
| 14^R^-R | GGATTGGGAATCCAAGCCAG | PCR-mapping assay |
| 15^R^-F | GGTGTATTCCCTTAGCATGC | PCR-mapping assay |
| 15^R^-R | GCTATCTATCTTTACCGCTT | PCR-mapping assay |
| 16^R^-F | AGATTTGGACACCAATATGG | PCR-mapping assay |
| 16^R^-R | GAGAATGGTAGATATGGCAG | PCR-mapping assay |
| 17^R^-F | CCTCGCTTGTAGTCTTCGTA | PCR-mapping assay |
| 17^R^-R | AAGGTGTTGGATCTGCTGAA | PCR-mapping assay |
| 18^R^-F | TAACCACCGTAGCTATATTG | PCR-mapping assay |
| 18^R^-R | GCGTAAAGTCACCACATTAT | PCR-mapping assay |
| 19^R^-F | CGGAGACCGTTAATAGAATG | PCR-mapping assay |
| 19^R^-R | CGGTAATGCAGAATATTTGG | PCR-mapping assay |
| 20^R^-F | TACCTCTTACTGGCATGCAT | PCR-mapping assay |
| 20^R^-R | CCAAGATGAAGCCAGTACTA | PCR-mapping assay |
| 21^R^-F | GTTGCAGCTTGAACTTGTTA | PCR-mapping assay |
| 21^R^-R | AGCATCAATTGGACATAGTG | PCR-mapping assay |
| RAP1-identify-F | CGCTGGAGCTACAAGTATTC | PCR Identification of *RAP1* gene |
| RAP1-identify-R | CTCATCAGCAGCGGTGTCAT | PCR Identification of *RAP1* gene |
| XVI-L-XY-KO-identify-F | TGACCTGACATGCACCAACA | PCR Identification of the deletion of X- and Y'-elements on XVI-L |
| XVI-L-XY-KO-identify-R | GCAGTGTAGATACCGTCCTT | PCR Identification of the deletion of X- and Y'-elements on XVI-L |
| XIII-L-X-KO-identify-F | CCGTATTAAAGAGGTGGCAC | PCR Identification of the deletion of the X-elements on XIII-L |
| XIII-L-X-KO-identify-R | CACACCAAGTTCTGATCCTC | PCR Identification of the deletion of the X-elements on XIII-L |
| I-L-X-KO-identify-F | CCGGATCACTAGCTCATTATC | PCR Identification of the deletion of the X-element on I-L |
| I-L-X-KO-identify-R | GCGGCTTGGAACATGTAGTA | PCR Identification of the deletion of the X-element on I-L |
| X-R-X-KO-identify-F | GGCGGTTTAGGCTTGTTCAT | PCR Identification of the deletion of the X-element on X-R |
| X-R-X-KO-identify-R | CCTCATCTTACCAGCTCACT | PCR Identification of the deletion of the X-element on X-R |
| XI-R-X-KO-identify-F | CTGTAGGCAGAGTGGCGAGT | PCR Identification of the deletion of the X-element on XI-R |
| XI-R-X-KO-identify-R | AAGGCCGAGAATAAGGGTAT | PCR Identification of the deletion of the X-element on XI-R |
| XIV-R-X-KO-identify-F | CTGTTGACCGGCACATTTGG | PCR Identification of the deletion of the X-element on XIV-R |
| XIV-R-X-KO-identify-R | CCTGAGTTGCGAATCTCACT | PCR Identification of the deletion of the X-element on XI-R |
| X-element-identify-F | GCACTTGCCTCAGCGGTCTATACCCT | PCR Identification of X-element |
| X-element-identify-R | ATTTTTATGTTTAGGTGATTTT | PCR Identification of X-element |
| Y’-element-identify-F | CTCACTAAGATAAGCGACTG | PCR Identification of Y’-element |
| Y’-element-identify-R | GTTAGACAAGGCCGTAGGGA | PCR Identification of Y’-element |
| X-R-MPH3-qPCR -F | TTTTTTTCATTATTGGCGGTTTG | qPCR |
| X-R-MPH3-qPCR -R | ACAGCCATTAAAAGAGAACCACTTC | qPCR |
| XVI-L-HSP32-qPCR -F | GGCGTTAGCGAGGATCAAGA | qPCR |
| XVI-L-HSP32-qPCR -R | TCCAGCAGATGCAAAGAATATTTT | qPCR |
| Y’-probe-F | CAGTTTAGCAGGCATCATCG | Y’-probe |
| Y’-probe-R | CGAGAACTTCAACGTTTGCC | Y’-probe |
| I-L-Type X-identify-R | GCTAACAGTGTATGTCCAATAAACAACAATGAAA | PCR amplification of the chromosome end sequence on I-L |
| X-R-Type X-identify-F | GGGCATCCATTACATTTCCGGTT | PCR amplification of the chromosome end sequence on X-R |
| XIII-L-Type X-identify-R | GGGCATCCATTACATTTCCGGTT | PCR amplification of the chromosome end sequence on XIII-L |
| XI-R-Type X-identify-F | AGATCGTAATTCATTACGTCAACATACCG | PCR amplification of the chromosome end sequence on XI-R |
| XVI-L-Type X-identify-R | CAAAACAAGTAGCAAGTCATAGCAAGAGA | PCR amplification of the chromosome end sequence on XVI-L |
| XIV-R-Type X-identify-F | GTATGAGTAGTGTCACATACAAGATTAAAGATGC | PCR amplification of the chromosome end sequence on XIV-R |
| TG primrer | CACACACCCACACACCACAC | complementary base pairing with telomere sequence |
| RAD52-identify-F | AAGGGAAACTAGGTTAAGAGCTAAAATATTG | PCR Identification of *RAD52* gene |
| RAD52-identify-R | TTCACCTTATCGATCTTTGCCAGAAAATCTTTATC | PCR Identification of *RAD52* gene |
| RAD51-identify-F | ACCTGTCATCGCAACGACGA | PCR Identification of *RAD51* gene |
| RAD51-identify-R | ACTTACCTGTCCTGAATTCA | PCR Identification of *RAD51* gene |
