## Supplementary file 2 for "Elimination of subtelomeric repeat sequences exerts little effect on telomere essential functions in *Saccharomyces cerevisiae*"

Supplementary file 2. The remaining subtelomeric elements in SY8 to SY13 strains

| Strain | Chromosome numbers | Number of X- element | Number of Y’- element | Location of the remaining X-element | Location of the remaining Y’-element |
| --- | --- | --- | --- | --- | --- |
| SY8 | 7 | 14 | 6 | Chr1 XIII-L XII-R  Chr2 VII-L VIII-R  Chr3 I-L II-R  Chr4 XVI-L XIV-R  Chr5 IX-L X-R  Chr6 III-L IV-R  Chr7 XV-L XI-R | Chr1 XIII-L XII-R  Chr2 VIII-R  Chr4 XVI-L  Chr5 IX-L  Chr6 IV-R |
| SY9 | 6 | 12 | 4 | Chr1 XIII-L XII-R  Chr2 VII-L X-R  Chr3 I-L II-R  Chr4 XVI-L XIV-R  Chr5 III-L IV-R  Chr6 XV-L XI-R | Chr1 XIII-L XII-R  Chr4 XVI-L  Chr5 IV-R |
| SY10 | 5 | 10 | 4 | Chr1 XIII-L XII-R  Chr2 VII-L X-R  Chr3 I-L IV-R  Chr4 XVI-L XIV-R  Chr5 XV-L XI-R | Chr1 XIII-L XII-R  Chr3 IV-R  Chr4 XVI-L |
| SY11 | 4 | 8 | 3 | Chr1 XIII-L XI-R  Chr2 VII-L X-R  Chr3 I-L IV-R  Chr4 XVI-L XIV-R | Chr1 XIII-L  Chr3 IV-R  Chr4 XVI-L |
| SY12 | 3 | 6 | 1 | Chr1 I-L X-R  Chr2 XIII-L XI-R  Chr3 XVI-L XIV-R | Chr3 XVI-L |
| SY13 | 2 | 4 | 2 | Chr1 I-L X-R  Chr4 XVI-L XI-R | Chr1 I-L  Chr4 XVI-L |

The remaining subtelomeric elements in SY8 to SY13 strains. The ‘Strain’ column lists the strain name. ‘Chromosome numbers’ indicates the chromosomes numbers in each strain. ‘Number of X-element’ and ‘Number of Y’-element’ shows the remaining subtelomere element in each strain. ‘Location of the remaining X-element’ and ‘Location of the remaining Y’-element’ lists their location; the Roman numerals, native chromosomes; the Arabic numerals, chromosome numbers of SYn stains. The information of each region is referred from the *S. cerevisiae* S288C genome (http://www.yeastgenome.org/).
