## Supplementary file 3 for "Elimination of subtelomeric repeat sequences exerts little effect on telomere essential functions in *Saccharomyces cerevisiae*"

| Supplementary file 3. Details of fusion points in­­ the “circular survivors” | | | | | |
| --- | --- | --- | --- | --- | --- |
| Strain | Chromosome | Deleted genetic elements | Retained subtelomeric elements | The length of the TG_1-3_ sequences at the junction points (bp) | The length of TG restriction fragments produced after XhoI digestion (kb) |
|  |  | Grey indicates genetic elements absent in SY12^YΔ^, SY12^XYΔ^ or SY12^XYΔ+Υ^ strain |  |  |  |
| SY12-*tlc1*Δ-C1 | I-L | *YAL067W-A*, *PAU8*, *ARS102*, *YAL068W-A*, *YAL069W* | - | ­- | - |
|  | X-R | *COS5*, *ARS1025*, *YJR162C* | - | - | - |
|  | XIII-L | *COS3* | - | - | - |
|  | XI-R | - | - | - | - |
|  | XVI-L | - | X | - | - |
|  | XIV-R | *COS10*, *YNR075C-A*, *PAU6*, *YNR077C* | - | - | - |
| SY12^YΔ^-*tlc1*Δ-C1 | I-L | *PAU8*, *ARS102*, *YAL068W-A*, *YAL069W* | - | - | - |
|  | X-R | *DAN1, ARS1022, DAN4*, *YJR151W-A*, *DAL5*, *PGU1*, *YJR154W*, *AAD10*, *THI11*, *ARS1023*, *YJR157W*, *HXT16*, *SOR1*, *ARS1024*, *MPH3*, *COS5*, *ARS1025*, *YJR162C* | - | - | - |
|  | XIII-L | - | X | 22 | 21.8 |
|  | XI-R | *SKG1*, *SIR1*, *ARS1123*, *FLO10*, *NFT1*, *YKR104W*, *VBA5*, *GEX2*, *YKRWomega1* | - |  |  |
|  | XVI-L | *PAU22*, *YPL283W-A*, *YPL283W-B*, *YRF1-7* | - | - | - |
|  | XIV-R | *COS10*, *YNR075C-A*, *PAU6*, *YNR077C* | - | - | - |
| SY12^XYΔ^-*tlc1*Δ-C1 | I-L | *YAL067W-A*, *PAU8*, *ARS102*, *YAL068W-A*, *YAL069W* | - | 34 | 25.2 |
|  | X-R | *RPS4A, YJR146W, HMS2, BAT2, YJR149W, DAN1, ARS1022, DAN4*, *YJR151W-A*, *DAL5*, *PGU1*, *YJR154W*, *AAD10*, *THI11*, *ARS1023*, *YJR157W*, *HXT16*, *SOR1*, *ARS1024*, *MPH3*, *COS5*, *ARS1025*, *YJR162C* | - |  |  |
|  | XIII-L | *COS3* | - | 13 | 18.0 |
|  | XI-R | *SKG1*, *SIR1*, *ARS1123*, *FLO10*, *NFT1*, *YKR104W*, *VBA5*, *GEX2*, *YKRWomega1* | - |  |  |
|  | XVI-L | *PAU22*, *YPL283W-A*, *YPL283W-B*, *YRF1-7* | - | - | - |
|  | XIV-R | *HXT17*, *MAN2*, *AIF1*, *COS10*, *YNR075C-A*, *PAU6*, *YNR077C* | - | - | - |
| SY12^XYΔ+Y^-*tlc1*Δ-C1 | I-L | *YAL067W-A*, *PAU8*, *ARS102*, *YAL068W-A*, *YAL069W* | - | 26 | 25.2 |
|  | X-R | *RPS4A, YJR146W, HMS2, BAT2, YJR149W, DAN1, ARS1022, DAN4*, *YJR151W-A*, *DAL5*, *PGU1*, *YJR154W*, *AAD10*, *THI11*, *ARS1023*, *YJR157W*, *HXT16*, *SOR1*, *ARS1024*, *MPH3*, *COS5*, *ARS1025*, *YJR162C* | - |  |  |
|  | XIII-L | *COS3* | - | 21 | 18.0 |
|  | XI-R | *SKG1*, *SIR1*, *ARS1123*, *FLO10*, *NFT1*, *YKR104W*, *VBA5*, *GEX2*, *YKRWomega1* | - |  |  |
|  | XVI-L | *ERR2*, *PAU22*, *YRF1-7* | - | - | - |
|  | XIV-R | *AIF1*, *COS10*, *YNR075C-A*, *PAU6*, *YNR077C* | - | - | - |
