## Supplementary file 4 for "Elimination of subtelomeric repeat sequences exerts little effect on telomere essential functions in *Saccharomyces cerevisiae*"

Supplementary file 4. Details of the creation of SY12 subtelomeric engineered strains

| Strain | Newly absent chromosome regions  (Distance from the telomere in bp) | Newly deleted/inserted subtelomeric elements | Remaining subtelomeric elements  (Chromosome ends) |
| --- | --- | --- | --- |
| SY12^YΔ^ | XVI-L: 1-9613 | Deletion of an X-element and a Y'-long element | Five X-elements  (XIII-L, I-L, X-R, XI-R, XIV-R) |
| SY12^YΔ+1XΔ^ | XIII-L: 1-3793 | Deletion of an X-element | Four X-elements  (I-L, X-R, XI-R, XIV-R) |
| SY12^YΔ+2XΔ^ | I-L: 1-3318 | Deletion of an X-element | Three X-elements  (X-R, XI-R, XIV-R) |
| SY12^YΔ+3XΔ^ | X-R: 1-4229 | Deletion of an X-element | Two X-elements  (XI-R, XIV-R) |
| SY12^YΔ+4XΔ^ | XI-R: 1-8477 | Deletion of an X-element | One X-element  (XIV-R) |
| SY12^XYΔ^ | XIV-R: 1-5591 | Deletion of an X-element | No X- or Y-elements |
| SY12^XYΔ+Y^ | XVI-L: 6655-10048 | Insertion of a Y'-long element | One Y’-long element  (XVI-L) |

Referred from the *S.cerevisiae* S288C genome (http://www.yeastgenome.org/).
