## Supplementary file 5 for "Elimination of subtelomeric repeat sequences exerts little effect on telomere essential functions in *Saccharomyces cerevisiae*"

Supplementary file 5. Quantitation of each survivor type in SY12 subtelomeric strains

| strain | I survivor | II survivor | Type X survivor | Uncharacterized survivor | Circular survivor | Total |
| --- | --- | --- | --- | --- | --- | --- |
| SY12 *tlc1*Δ | 8 | 1 | 12 | 19 | 10 | 50 |
| SY12^YΔ^ *tlc1*Δ | 0 | 2 | 0 | 22 | 26 | 50 |
| SY12^XYΔ^ *tlc1*Δ | 0 | 4 | 0 | 22 | 24 | 50 |
| SY12^XYΔ+Y^ *tlc1*Δ | 0 | 7 | 0 | 21 | 22 | 50 |
| SY12^XYΔ^ *tlc1*Δ *rad51*Δ | 0 | 8 | 0 | 8 | 9 | 25 |
