## Supplementary file6 for "Elimination of subtelomeric repeat sequences exerts little effect on telomere essential functions in *Saccharomyces cerevisiae*"

| Supplementary file 6. Yeast strains used in this study | | |
| --- | --- | --- |
| Strain | Genotype | Source |
| BY4742 | *MATα his3*Δ*1 leu2*Δ*0 lys2*Δ*0 ura3*Δ*0* | Euroscarf |
| SY12 | *MATα his3*Δ*1 leu2*Δ*0 lys2*Δ*0 ura3*Δ*0* | (Shao et al., 2018) |
| WZJ0044 | BY4742 *tlc1*Δ::*HIS3/CEN* pRS316-*TLC1* | This study |
| WZJ0045 | SY1 *tlc1*Δ:: *HIS3/CEN* pRS316-*TLC1* | This study |
| WZJ0046 | SY3 *tlc1*Δ::*HIS3/CEN* pRS316-*TLC1* | This study |
| WZJ0047 | SY5 *tlc1*Δ::*HIS3/CEN* pRS316-*TLC1* | This study |
| WZJ0048 | SY7 *tlc1*Δ::*HIS3/CEN* pRS316-*TLC1* | This study |
| WZJ0049 | SY8 *tlc1*Δ::*HIS3/CEN* pRS316-*TLC1* | This study |
| WZJ0050 | SY9 *tlc1*Δ::*HIS3/CEN* pRS316-*TLC1* | This study |
| WZJ0051 | SY10 *tlc1*Δ::*HIS3/CEN* pRS316-*TLC1* | This study |
| WZJ0052 | SY11 *tlc1*Δ::*HIS3/CEN* pRS316-*TLC1* | This study |
| WZJ0053 | SY12 *tlc1*Δ::*HIS3/CEN* pRS316-*TLC1* | This study |
| WZJ0054 | SY12^YΔ^ | This study |
| WZJ0055 | SY12^YΔ+1XΔ^ | This study |
| WZJ0056 | SY12^YΔ+2XΔ^ | This study |
| WZJ0057 | SY12^YΔ+3XΔ^ | This study |
| WZJ0058 | SY12^YΔ+4XΔ^ | This study |
| WZJ0059 | SY12^XYΔ^ | This study |
| WZJ0060 | SY12^XYΔ+Y^ | This study |
| WZJ0061 | BY4742 *sir2*Δ::*HIS3* | This study |
| WZJ0062 | SY12 *sir2*Δ::*HIS3* | This study |
| WZJ0063 | SY12^YΔ^ *sir2*Δ::*HIS3* | This study |
| WZJ0064 | SY12^XYΔ^ *sir2*Δ::*HIS3* | This study |
| WZJ0065 | SY12^XYΔ+Y^ *sir2*Δ::*HIS3* | This study |
| WZJ0066 | SY12^YΔ^ *tlc1*Δ::*HIS3/CEN* pRS316-*TLC1* | This study |
| WZJ0067 | SY12^XYΔ^ *tlc1*Δ::*HIS3/CEN* pRS316-*TLC1* | This study |
| WZJ0068 | SY12^XYΔ+Y^ *tlc1*Δ::*HIS3/CEN* pRS316-*TLC1* | This study |
| WZJ0069 | SY12 *tlc1*Δ::*HIS3 yku70Δ::LEU2/CEN* pRS316-*TLC1* | This study |
| WZJ0070 | SY12^YΔ^ *tlc1*Δ::*HIS3 yku70Δ::LEU2/CEN* pRS316-*TLC1* | This study |
| WZJ0071 | SY12^XYΔ^ *tlc1*Δ::*HIS3 yku70Δ::LEU2/CEN* pRS316-*TLC1* | This study |
| WZJ0072 | SY12^XYΔ+Y^ *tlc1*Δ::*HIS3 yku70Δ::LEU2/CEN* pRS316-*TLC1* | This study |
| WZJ0073 | SY12 *tlc1*Δ::*HIS3 rad51Δ::LEU2/CEN* pRS316-*TLC1* | This study |
| WZJ0074 | SY12 *tlc1*Δ::*HIS3 rad52Δ::LEU2/CEN* pRS316-*TLC1* | This study |
| WZJ0075 | SY12^YΔ^ *tlc1*Δ::*HIS3 rad52Δ::LEU2/CEN* pRS316-*TLC1* | This study |
| WZJ0076 | SY12^XYΔ^ *tlc1*Δ::*HIS3 rad52Δ::LEU2/CEN* pRS316-*TLC1* | This study |
| WZJ0077 | SY12^XYΔ+Y^ *tlc1*Δ::*HIS3 rad52Δ::LEU2/CEN* pRS316-*TLC1* | This study |
| WZJ0078 | SY12XYΔ tlc1Δ::HIS3 rad50Δ::LEU2/CEN pRS316-TLC1 | This study |
| WZJ0079 | SY12XYΔ tlc1Δ::HIS3 rad51Δ::LEU2/CEN pRS316-TLC1 | This study |
